# Temporal placement of RNA modifications under 5-fluorouracil treatment in human cell culture

**DOI:** 10.1101/2024.10.21.619086

**Authors:** Maximilian Berg, Chengkang Li, Stefanie Kaiser

## Abstract

Recent studies have explored the dynamic changes in RNA modification across various organisms, often in response to stressors like chemical agents or drugs. Among these, the anticancer drug 5-fluorouracil (5-FU) has been evaluated regarding its impact on RNA and DNA. Although different alterations in RNA modification have been associated with 5-FU, a mechanistic insight into the dynamic effects on RNA modification is missing. In this study, we provide a detailed insight into the dynamic modification changes for tRNA and rRNA of HEK293T cells exposed to 5-FU using Nucleic Acid Isotope Labeling coupled Mass Spectrometry (NAIL-MS). Consistent with existing literature, we observed a constant loss of m^5^U in newly transcribed tRNA during 5-FU exposure. In addition to m^5^U, we identified a loss of m^5^Um and an increase of Um within tRNA^Lys^_UUU_. While cells recover from a 24-hour 5-FU treatment, we found changed abundances of modifications like an increase of mcm^5^U and m^5^C and a decrease of m^7^G and m^1^A in total tRNA. Moreover, we demonstrate that 5-FU affects various rRNA subtypes, altering both their modification stoichiometry and abundance. Taken together, we use NAIL-MS to deconvolute RNA modification abundance changes caused by drug treatment and we uncover patterns of RNA modification adaptation.

## INTRODUCTION

RNA as a key player in cellular biology links genetic information encoded in DNA with the phenotypic information expressed as proteins (1). To accurately fulfil this purpose, the four canonical RNA nucleosides cytidine, uridine, adenosine and guanosine undergo additional modification. These modifications, implemented by writer enzymes, encompass a diverse chemical variety, including methylation, thiolation and other group additions (2). The landscape of RNA modifications is evidenced by the identification of over 150 different modifications (3) to date, with ongoing discoveries underlined by recently published literature (4–6).

Among RNA species, tRNA exhibits the highest density and diversity of modifications (2), which contributes to both structural stability and biological function (7). For instance, m^5^U is a commonly found tRNA modification implemented by TRMT2A, affecting the secondary tRNA structure as well as translational fidelity (8,9). Notably, modifications within the anticodon loop of tRNA directly influence the translation process, particularly at the wobble position 34 (10). The presence of modified residues at this position can modulate codon-anticodon interactions, thereby impacting translational fidelity and efficiency (11). The interplay between tRNA modifications and codon-anticodon pairing led to further investigations in stress response studies. Early studies revealed that certain stressors cause alterations in the tRNA modification landscape, a phenomenon termed tRNA modification reprogramming (12). Recent research highlighted the association between adaptive stress responses to environmental stress, including hypoxia and chemical toxins and specific modifications, such as wobble uridine modifications like mcm^5^U and mcm^5^s^2^U (13,14). These findings underscore the significance of tRNA modifications in cellular homeostasis and stress response.

The modification landscape of rRNA predominantly consists of uridine isomerizations (Ψ) and 2′-O-methylations (Nm) (3,15,16). Most of these modifications are introduced early during rRNA maturation in the nucleolus, primarily guided by small nucleolar RNAs (snoRNAs) (17). In addition to ribose-methylated nucleosides, other modifications, such as m^3^U, m^5^C, ac^4^C, and m^6,6^A, are found in 28S- and 18S-rRNA (3). While most Nm modifications are known to be introduced during the chromatin-associated stage, either prior to or during rRNA processing, the precise timing for many other modifications remains unclear (18). Similar to tRNA, several studies have reported stress-induced reprogramming on rRNA (19) or ribosome level, e.g. p53-mediated ribosomal stress (20).

Of particular interest is the stress response caused by pharmacological agents. Often, the mechanism of action of commonly used drugs is known and their effect is elucidated on the cellular level. However, how drugs potentially interfere with RNA modifications remains mostly elusive, even though there is evidence that certain drugs disrupt RNA modification patterns. Examples of known effects on RNA modifications are given for nucleoside- and nucleobase-mimicking drugs, such as 5-fluorouracil (5-FU) and others (21,22). As described for 5-FU, the mechanism of action is inhibition of thymidylate synthase (23), which leads to DNA damage and cell cycle arrest (24). Despite this, 5-FU is incorporated into nucleic acids and interferes with different enzymes of nucleic acid metabolism (25). For example, writer enzymes like PUS1 and TRMT2A are inhibited under 5-FU treatment, which results in aberrant absolute modification abundances for pseudouridine (Ψ) and 5-methyluridine (m^5^U) (26,27). Furthermore, 5-FU impairs rRNA processing and maturation, making the effects on RNA level even more diverse (28,29). Intrigued by the impact of 5-FU on RNA modification, we wanted to specifically investigate its effects on tRNA and rRNA modifications and cellular homeostasis in human cell lines. Using nucleic acid isotope labeling coupled mass spectrometry (NAIL-MS) (30), we elucidated the dynamic changes in tRNA modification profiles induced by 5-FU treatment. Together with tRNA isoacceptor and proteomic data, we show that there is a complex interplay between RNA modification abundances and RNA abundance in tRNA and rRNA under 5-FU treatment. Altogether, our work sheds light on the broader implications of RNA dysregulation in cellular responses to pharmacological interventions.

## MATERIAL AND METHODS

### Chemicals and Reagents

All chemicals and reagents were purchased from Sigma Aldrich (St. Louis, MO, USA) unless stated otherwise. ^13^C_5_,^15^N_2_-Uridine and ^15^N_5_-Adenine were obtained from Cambridge Isotope Laboratories (Tewksbury, MA, USA). Nucleoside standards Pseudouridine (Ψ), 1-methyladenosine(m^1^A), N3-methylcytidine (m^3^C), N7-methylguanosine (m^7^G), 5-methylcytidine (m^5^C), 5-methyluridine (m^5^U), 2’-O-methylcytidine (Cm), 2’-O-methylguanosine (Gm), 2’-O-methyladenosine (Am), 2’-O-methyluridine (Um), 1-methylguanosine (m^1^G), N2-methylguanosine (m^2^G), N2,N2-dimethylguanosine (m^22^G), inosine (I), 5-carbamoylmethyluridine (ncm^5^U), N4-acetylcytidine (ac^4^C), N6-metyhladenosine (m^6^A) and 5-methoxycarbonylmethyl-2-thiouridine (mcm^5^s^2^U) were obtained from Carbosynth (Newbury, UK). N6-threonylcarbamoyladenosine (t^6^A), N6,N6-dimethyladenosine (m^66^A) and 5-methyl-2-O’-dimethyluridine (m^5^Um) were obtained from TRC (New York, Canada). N3-methyluridine (m^3^U), N6-isopentenyladenosine (i^6^A), queuosine (Q) and 5-methoxycarbonylmethyluridine (mcm^5^U) were generous gifts from the Dedon lab. 1-Methylinosine (m^1^I) was a generous gift from STORM Therapeutics LTD (Cambridge, UK).

### Cell culture

Standard growth medium for HEK293 culture was Dulbecco’s modified Eagle’s medium (DMEM) D6546 high glucose supplemented with 10% FBS, 0.584 g L^-1^ L-glutamine and 8 µg L^-1^ queuine. Cells were split 1:7 every 2 to 3 days to counter overgrowth. Cells were incubated at 37 °C and 10% CO_2_ for pH adjustment, provided by a CellXpert C170i (Eppendorf, Hamburg, Germany). To prevent contamination, sterile consumables were used and work was performed in a laminar air flow hood (HeraSafe 2025, Thermo Fisher Scientific, Waltham, MA, USA). DMEM D0422 without cysteine and methionine was used for all experiments which included nucleic acid isotope labeling. DMEM D0422 was supplemented with 10% dialyzed FBS, 0.584 g L^-1^ glutamine, 0.063 g L^-1^ cysteine, 0.03 g L^-1^ methionine, 0.05 g L^-1^ uridine, 0.015 g L^-1^ adenine and 8 µg L^-1^ queuine. Methionine, uridine and adenine were either added as unlabeled or labeled compounds depending on the desired labeling. For drug incubation, 5-FU (100 µM) and/or actinomycin D (1 µg mL^-1^) were added to the respective media.

### Cell lysis for RNA isolation

Cells were directly harvested in cell culture dishes using 1 mL TRI reagent for T25 flasks or 0.5 mL TRI for smaller dishes. Total RNA was isolated according to the manufacturers protocol. The aqueous phase was mixed with isopropanol in a 1:1 ratio to precipitate total HEK RNA. After overnight incubation at -20 °C, samples were centrifuged at 12.000 g for 30 min at 4 °C. The supernatant was removed and RNA pellets were washed twice with 180 µL 70 % ethanol and centrifuged at 12.000 g for 10 min at 4 °C. After removing the supernatant, the samples were placed on the bench for 10 min to let the remaining ethanol evaporate. RNA was reconstituted in 30 µL water.

### Cell lysis for protein isolation

Cells were dissociated using 1 mL of Gibco™ TrypLE™ Express Enzyme (1X) (Thermo Fisher Scientific, Waltham, MA, USA) for 2 min at 37°C. The reaction was quenched using 4 mL of DMEM D6546 growth medium and the cells were centrifuged at 1.300 g for 3 min at room temperature. Subsequently, the supernatant was removed, and the cells were washed with 1 mL ice-cold PBS without calcium and magnesium (1X) twice and resuspended with another 1 mL PBS. An aliquot of 1 Mio cells was transferred into a new tube and centrifuged at 1.300 g for 3 min at room temperature, which was then used for cell lysis. After removing the supernatant, the cell pellets were re-suspended in 200 µL ice-cold protein extraction buffer [4 M Urea/50 mM Tris (pH 8.0) + 0.1 % *n*-Dodecyl-β-D-Maltoside (DDM) + Halt™ Protease and Phosphatase Inhibitor Single-Use Cocktail (1X) (Thermo Fisher Scientific, Waltham, MA, USA) + 5 mM EDTA]. Samples were then sonicated using an Ultrasonic processor UP200St coupled with a VialTweeter (Hielscher, Teltow, Germany) at an amplitude of 60%, a pulse cycle of 60% and a duration of 20s for 5 times with intermediate cooling on ice. The lysates were centrifuged at 15.000 g for 30 min at 4 °C and the supernatant containing the protein fraction was used for further processing for proteomic analysis as described in the “**Sample preparation for LC-MS/MS-based Proteomics**” section below.

### tRNA and rRNA isolation

tRNA was purified using size exclusion chromatography or 10% TBE-urea-PAGE.

#### SEC

SEC was performed on an Agilent HPLC 1100 Series (Agilent Technologies, Santa Clara, CA, USA) equipped with an AdvanceBio SEC 300 Å, 2.7 µm, 7.8 x 300 mm for tRNA purification and an AdvanceBio SEC 1000 Å, 2.7 µm, 7.8 x 300 mm for rRNA purification using 0.1 M ammonium acetate buffer (pH 7) at a flow rate of 1 mL min^-1^ and a temperature of 40 °C. The excess solvent was reduced to 50 µL using a Savant SpeedVac SPD120 (Thermo Fisher Scientific, Waltham, MA, USA) an RNA was precipitated using 0.1x vol. 5 M ammonium acetate and 2.5x vol. absolute ethanol. After overnight incubation at -20 °C, samples were centrifuged at 12.000 g for 30 min at 4 °C. The supernatant was removed and RNA pellets were washed with 180 µL 70 % ethanol and centrifuged at previous conditions for 10 min. Supernatant was removed again and samples were placed on the bench for 10 min to let the remaining ethanol evaporate. RNA was reconstituted in 30 µL of water.

#### PAGE

For tRNA purification via polyacrylamide gel electrophoresis, total RNA was separated by 10% TBE-urea PAGE in 1X TBE buffer (reagents obtained from Carl Roth, Karlsruhe, Germany). Samples were mixed 1:1 with formamide loading buffer, denatured at 90 °C for 2 min and an aliquot of 10 µg RNA was immediately loaded on the gel. The gel was run for 40 min at constant 275 V. Dyeing of tRNA was done with Gelstain Red (Carl Roth GmbH, Karlsruhe, Germany) and visualized using a ChemiDoc MP Imaging System (Bio-Rad Laboratories GmbH, Feldkirchen, Germany). tRNA bands were cut out using a scalpel and crushed with the blasé of the scalpel. The crushed tRNA bands were eluted in a total volume of 300 µL 0.5 M ammonium acetate and subsequently eluted a second time using 150 µL ammonium acetate. Both elusion steps were combined and filtered using 0.45 µm Nylon membranes at 6.000 g for 8 min at room temperature. The excess solvent was reduced to 50 µL using a Savant SpeedVac SPD120 (Thermo Fisher Scientific, Waltham, MA, USA) and tRNA was precipitated using 0.1x vol. 5 M ammonium acetate and 2.5x vol. absolute ethanol. After overnight incubation at -20 °C, samples were centrifuged at 12.000 g for 30 min at 4 °C. The supernatant was removed and RNA pellets were washed with 180 µL 70 % ethanol and centrifuged at previous conditions for 10 min. Supernatant was removed again and samples were placed on the bench for 10 min to let the remaining ethanol evaporate. RNA was reconstituted in 30 µL of water.

### tRNA isoacceptor and tRNA fragment purification

For tRNA-isoacceptor purification, 1 µg of total tRNA was mixed with 100 pmol reverse complementary, biotinylated DNA oligonucleotide in a total volume of 100 µL 5X SSC buffer (0.75 M NaCl, 75 mM trisodium citrate pH 7). The mixture was incubated at 90 °C for 3 min for denaturation followed by a hybridization step at 65 °C for 10 min. For each sample, 25 µL Magnetic Dynabeads® Myone^TM^ Streptavidin T1 (Thermo Fisher Scientific, Waltham, MA, USA) were primed three times using Bind and Wash buffer (B&W, 5 mM Tris-HCl pH 7.5, 0.5 M EDTA, 1 M NaCl) and once using 5X SSC buffer. An aliquot of 25 µL magnetic beads in 5X SSC buffer was added to each sample and incubated at RT for 30 min at 600 rpm. Magnetic racks were used to separate the Beads from unbound tRNA and the magnetic beads were washed once using 50 µL 1X SSC buffer and three times using 25 µL 0.1 X SSC buffer. Elution of the desired tRNA was carried out in 20 µL water at 75 °C for 3 min. Samples were directly used for LC-MS preparations.

For fragment analysis of tRNA^Lys^_UUU_, 1 µg of total tRNA was mixed with 100 pmol reverse complementary, biotinylated DNA oligonucleotide in a total volume of 45 µL 1X RNase T1 buffer (25 mM Tris-HCl (pH = 7.5), 100 mM NaCl). The mixture was incubated at 90 °C for 3 min for denaturation followed by a hybridization step at 65 °C for 10 min. Subsequently, 5U of RNase T1 (NEB, Ipswich, USA) were added for a total volume of 50 µL and incubated for 1 h at 37 °C. After digestion, the samples were supplied with SSC buffer for a final volume of 100 µL 5X SSC buffer. Further procedures after adding the magnetic dynabeads were carried out as described above.

### Northern Blotting

Total RNA was separated by 12% TBE-urea PAGE in 1X TBE buffer. Samples were mixed 1:1 with formamide loading buffer, denatured at 90 °C for 2 min and an aliquot of 10 µg RNA was immediately loaded on the gel. The gel was run for 40 min at constant 275 V and the RNA was subsequently transferred onto a Hybond-N^+^ nylon membrane (GE healthcare, Chicago, USA) at 1.5 A with a Trans-Blot Turbo Transfer System (Bio-Rad Laboratories GmbH, Feldkirchen, Germany) for 7 min. The RNA was crosslinked with UV light at 120 mJ cm^-2^ and the nylon membrane was subsequently incubated for 30 min in hybridization buffer (5X Denhardt’s solution, 1% SDS, 6.6X SSPE buffer). 100 pmol of the respective 3’ and 5’ Cyanine-3 modified oligonucleotide probe was added and incubated over night at 37°C in a shaking incubator with a shaking oscillation of 200 rpm. The next day the nylon membrane was washed for 10 min with washing buffer (0.5% SDS in 2X SSPE buffer) at 200 rpm and imaged using a ChemiDoc MP Imaging System (Bio-Rad Laboratories GmbH, Feldkirchen, Germany). The signals of the respective isoacceptor were normalized using 5S-rRNA as loading control.

### Preparations for nucleoside LC-MS/MS

Total rRNA was diluted to a final concentration of 15 ng µL^-1^ and a final volume of 20 µL. Total tRNA was diluted to a final concentration of 10 ng µL^-1^ and a final volume of 20 µL. Single tRNA isoacceptor samples were used as described in the respective methods section. RNA was then digested to single nucleosides using a fresh prepared digestion master mix containing 2 U benzonase, 2 U alkaline phosphatase and 0.2 U phosphodiesterase I in 5 mM Tris (pH 8) and 1 mM MgCl_2_ containing buffer. To avoid deamination and oxidation of nucleosides, 0.5 µg of pyrimidine deamination inhibitor tetrahydrouridine, 0.1 µg of purine deamination inhibitor pentostatin and 1 µM of antioxidant butylated hydroxytoluene were added. The digestion mixture in a total volume of 35 µL was incubated for 2h at 37 °C and 10 µL of LC-MS buffer was added afterwards. For quantitative analysis, a calibration mixture was prepared using synthetic nucleosides. The calibration solutions ranged from 0.025 to 100 pmol for canonical nucleosides and from 0.00125 pmol to 5 pmol for modified nucleosides, except of pseudouridine which ranged from 0.005 to 20 pmol. 10 µL of each sample and calibration was injected into the LC-MS system for analysis. Additionally, 1 µL of previously prepared and digested stable isotope labeled internal standard (31) was co-injected.

### LC-MS/MS of Nucleosides

For quantitative mass spectrometry of nucleosides, an Agilent 1290 Infinity II equipped with a diode-array detector (DAD) combined with an Agilent Technologies G6470A Triple Quad system and electrospray ionization (ESI-MS, Agilent Jetstream) was used. Operating parameters: positive-ion mode, skimmer voltage of 15 V, cell accelerator voltage of 5 V, N_2_ gas temperature of 230 °C and N_2_ gas flow of 6 L/min, N_2_ sheath gas temperature of 400 °C with a flow of 12 L/min, capillary voltage of 2500 V, nozzle voltage of 0 V, nebulizer at 40 psi. The instrument was operated in dynamic multiple reaction monitoring mode (dMRM).

For separation a Synergi, 2.5 µm Fusion-RP, 100 Å, 100 x 2 mm column (Phenomenex, Torrance, California, USA) at 35 °C and a flow rate of 0.35 mL/min was used. Mobile phase A consisted of 5 mM aqueous NH_4_OAc buffer, brought to a pH of 5.3 with glacial acetic acid (65 µL/L) and mobile phase B consisted of organic solvent acetonitrile (Roth, Ultra-LC-MS grade). The gradient started at 100 % A for 1 min and an increase of 10% B over a period of 4 min afterwards. B was then increased to 40 % for 2 min and maintained for 1 min before returning to 100% A over a period of 0.5 min followed by a re-equilibration period for 2.5 min.

### Data analysis of nucleoside LC-MS/MS

Raw data was analyzed using quantitative and qualitative MassHunter Software from Agilent. The signals for each nucleoside from dMRM acquisition were integrated along with the respective SILIS. The signal areas of nucleoside and respective SILIS were set into relation to calculate the nucleoside isotope factor (NIF), eq. 1:

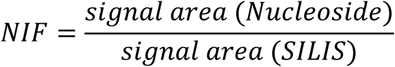

The nucleoside isotope factor was then plotted against the molar amount of each calibration and regression curves were plotted through the data points. The slopes represent the respective relative response factors for the nucleosides (rRFN) and enable absolute quantification of nucleosides. Calibration curves were plotted automatically by quantitative MassHunter software from Agilent. Molar amounts of nucleosides in samples were calculated using the signal areas of the target compounds and SILIS in the samples and the respective rRFN, determined by calibration measurements. This step was also done automatically by quantitative MassHunter software. The detailed calculation is depicted in eq. 2:

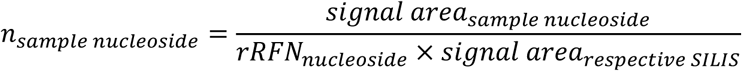

The molar amount of modified nucleosides was then normalized to respective tRNA population to calculate the amount of modification per tRNA. This was done using the expected amount of canonical nucleosides (taken from databases and/or sequencing data) of the respective tRNA population (eq. 3):

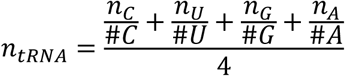

The molar amount of modified rRNA nucleosides was normalized to the molar amount of 1000 canonical nucleosides of the respective rRNA-type (eq. 4):

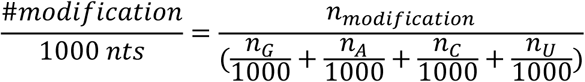

For experiments including nucleic acid isotope labeling, the isotopologues were normalized to the labeled canonical nucleosides to differentiate between pre-existing modifications and new modifications in the respective tRNA transcripts.

Statistical analysis was done using Excel and Graphpad Prism. Statistical significance was calculated by Welch’s t-test (p < 0.05 was considered significant).

### Sample preparation for LC-MS/MS-based Proteomics

Protein extracts of samples were reduced using Tris(2-carboxyethyl)phosphinehydrochloride (TCEP, final conc. 15 mM) and alkylated (via carbamidomethylation) using chloroacetamide (CAA, final conc. 40 mM) with both incubations being performed for 30 min at 30 °C in the dark. Samples were diluted 10 times with 50 mM ammonium bicarbonate (pH 8.0) to lower the urea and DDM concentration for maximized protease activity. Pierce™ trypsin protease (Thermo Scientific, Rockford, IL, USA) was added to each sample to reach 2 % (w/w) trypsin to protein ratio and incubated overnight at 37 °C. The detergent (i.e. DDM) within the protein digests was removed using water-saturated ethyl acetate as described in previously (32). Afterwards, the DDM-free peptide samples were desalted with 0.5 % formic acid and eluted with 0.5 % formic acid/80 % acetonitrile using SepPak C18 cartridges (Waters, Milford, MA, USA) with the help of CHROMABOND® SPE vacuum manifold (Macherey-Nagel GmbH & Co. KG, Duren, Germany). Samples were then dried in a Savant SpeedVac SPD120 (Thermo Fisher Scientific, Waltham, MA, USA) and resuspened in 2 % formic acid before LC-MS/MS analysis.

### LC-MS/MS-based Proteomics

Proteomic mass spectrometry measurements were performed on an Orbitrap Q Exactive plus (Thermo Fisher Scientific, Waltham, MA, USA) coupled to an UltiMate™ 3000 Nano-HPLC via Nanospray Flex ion source (Thermo Fisher Scientific, Waltham, MA, USA). Each peptide sample was first loaded on a PepMap™ Neo Trap Cartridge (Thermo Fisher Scientific, Waltham, MA, USA) at 5 μL/min for 5 min using 100% mobile phase A and subsequently reverse eluted onto an Acclaim™ PepMap™ 100 C18 analytical column (150 mm x 0.075 mm, 2 µm, 100 Å) at a flow rate of 0.3 μL/min. Mobile phase A consisted of 0.1% formic acid and mobile phase B consisted of 0.1 % formic acid/acetonitrile. The gradient started with 5% B and was then increased to 35 % over a period of 95 min. B was then further increased to 99% over a period of 10 min and maintained for 30 min. Afterwards, B was decreased to 0% over 5 min and maintained for 10 min. The mobile phase composition was then equilibrated to the initial condition (i.e. 5% B) in 1 min and maintained until the end of the run for 4 min. The temperature of the column oven was set to be 30 °C. The mass spectrometer was operated in Full MS/data dependent MS^2^ (dd-MS^2^) mode. The parameters were as followed: Polarity: positive; Chrom. Peak width (FWHM): 15 s; Full MS microscans: 1; Full MS resolution: 70 k; Full MS AGC target: 1e6; Full MS maximum injection time: 50 ms; Full MS number of scan ranges: 1; Full MS scan range: 375 – 1500 m/z; dd-MS^2^ resolution: 35 k; dd-MS^2^ AGC target: 2e4; dd-MS^2^ maximum injection time: 35 ms; dd-MS^2^ loop count: 15; dd-MS^2^ MSX count: 1; dd-MS^2^ Isolation window: 1.7 m/z; dd-MS^2^ Isolation offset: 0.0 m/z; dd-MS^2^ normalized collision energy (NCE): 10; dd-MS^2^ spectrum data type: profile. Data dependent settings: Minimum AGC target: 1e4; apex trigger: 2 – 15 s; charge exlusion: unassigned, >8; peptide match: preferred; exclude isotopes: on; dynamic exlusion: 30 s.

### Proteomic data evaluation

Proteomic data obtained by MS measurement was analyzed using MaxQuant version 1.6.5.0 and Perseus version 1.6.15.0 software. MS signals were annotated using a *H. sapiens* fasta-file from NCBI and quantified. Data points originating from potential contaminants were excluded and missing values were replaced from a normal distribution using Perseus’ default settings (width 0.3, down shift 1.8). The significance line was calculated using Perseus’ default settings (t-test, 250 randomizations, FDR 0.05, S0 0.1). Mass spectrometry proteomics data has been deposited to the ProteomeXchange Consortium via PRIDE (33) partner repository with the dataset identifier PXD051672 and 10.6019/PXD051672.

### Codon usage analysis and isoacceptor usage analysis

Human cDNA sequences, which represented the filtered transcript sequence from start to stop codon for proteins of interest (up- and downregulated proteins from proteomics), were provided from GenBank. The GSCU and isoacceptor usage of all human proteins and all higher-abundant proteins were calculated as recently described (14). For statistics student t-test was performed between all human proteins and the higher-abundant proteins.

## RESULTS

### 5-FU causes changes in tRNA modification abundance

To assess which tRNA modification abundances change during acute and after 5-FU treatment, we treated HEK 293T cells with various doses of 5-FU to determine a sub-lethal dose. At 100 µM 5-FU, approximately 50 % of the cells survived 24h of treatment (Figure S1). At this survival rate, we expect to see all major tRNA modification adaptations while still having enough cells to work with. Then, we exposed HEK cells to 100 µM 5-FU for either 6, 12 or 24 hours to assess tRNA modification abundances during acute exposure. In the next experimental setup, we treated cells for 24 hours with 5-FU and replaced the 5-FU supplied medium with fresh medium. Through this medium exchange, we wanted to further determine 5-FU effects after drug removal and harvested the cells after 6h and 12h medium exchange respectively (i.e. 30 and 36 hours after experiment initiation). Total RNA from all samples was harvested and total tRNA was purified using either polyacrylamide gel electrophoresis (PAGE) or size exclusion chromatography. Exemplary gels, chromatograms and quality controls of purified total tRNA by chip gel electrophoresis are shown in Figure S2. Total tRNA was hydrolysed to nucleosides and subjected to quantification using our established isotope-dilution protocol. Figure S3 shows that one of the active metabolites, 5-fluorouridine, was incorporated into tRNA in a time-dependent manner. We find 0.7 mol 5-FU (detected as 5-fluoruridine) per mol total tRNA after 24 hours of exposure to 100 µM 5-FU (Figure S3). Figure 1A shows the change in tRNA modification abundance of each time point in comparison to the starting point before 5-FU exposure.

**Figure 1.**
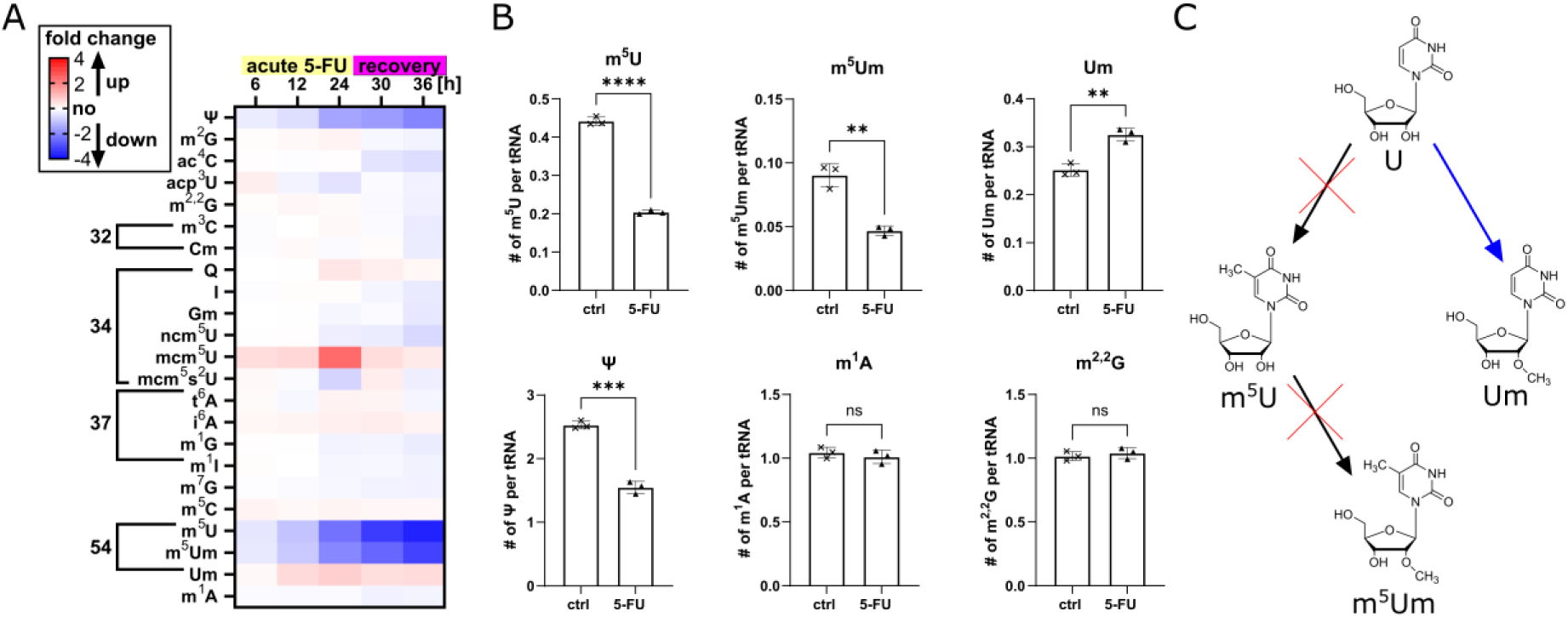
Relative and absolute abundance of total tRNA modifications during exposure to 5-fluoruracil (5-FU). **(A)** Heatmap showing the fold change in HEK293T total tRNA modification abundances under acute 5-FU exposure (6h, 12h and 24h) and after 24h 5-FU exposure following recovery (30h and 36h). Colour code: Red: higher abundance; white: no abundance change; blue: lower abundance. As the location of many tRNA modifications is limited to a certain position, we have ordered the modifications in ascending, expected order. **(B)** Selected modifications and their absolute abundance in total tRNA after 24h 5-FU exposure. **(C)** Modification pathway of C5-methylated uridine modifications and contributions of 5-FU to modification changes. (Statistics: Welchs t-test, p > 0.05 (ns), p < 0.05 (*), p < 0.01 (**), p < 0.001 (***), p < 0.0001 (****)).

Most modifications, such as N2-methylguanosine (m^2^G), N2,N2-dimethylguanosine (m^22^G), inosine (I), 5-methylcytidine (m^5^C) or 1-methyladenosine (m^1^A) appear to have a constant abundance throughout the experiment. Other modifications, like 5-methyluridine (m^5^U) or pseudouridine (Ψ), show a clear and exposure-time dependent decrease in abundance as expected from previous studies (22,34). This effect is related to the mode of action of 5-FU, which covalently inhibits enzymes such as TRMT2 (m^5^U writer) and PUS (Ψ writer) (35,36). From the modomics database, we know that m^5^U is found at position 54 in at least 8 out of 47 human cytosolic tRNAs. In tRNA^Lys^_UUU_ the ribose-methylated variant of m^5^U (m^5^Um) was reported (37). The absolute values for m^5^U and m^5^Um reflect the occurrence in total tRNA, respectively, as 0.44±0.01 mol m^5^U and only 0.09±0.01 mol m^5^Um is found per mol total tRNA (Figure 1B, for absolute values of other modifications see Figure S4). After 24 hours of 5-FU exposure, the amounts of m^5^U and m^5^Um decrease by 0.2 mol and 0.05 mol, respectively. Interestingly, the third modification of the m^5^Um pathway (Figure 1C), namely 2’-O-methyluridine (Um), increases by 0.05 mol per tRNA during exposure. This indicates that m^5^U formation through TRMT2A is disrupted, whereas the writer for m^5^Um is not affected by 5-FU. Therefore, U54 in the respective isoacceptors will be ribose methylated, which results in the formation of Um instead of m^5^Um (Figure 1C).

As 5-FU inhibits C5 methylation of uridine, we were interested in the effects on other C5-modified uridine derivatives. These are uniquely found at position 34 of tRNA, they are chemically highly diverse, and they alter translation (38,39). In the acute phase of exposure, we observed a time-dependent increase for mcm^5^U, but not for mcm^5^s^2^U and ncm^5^U. This indicates that the ELP complex is not inhibited by 5-FU. In the recovery phase, all three modifications showed changed abundances. Although this data is received through state-of-the-art LC-MS/MS analysis, biological interpretation is limited due to the complex nature of the analysed sample. This limitation arises through total tRNA being a mixture of 47 cytosolic tRNA isoacceptors and a mixture of tRNAs present before 5-FU exposure and new tRNAs transcribed during 5-FU exposure. Therefore, it is difficult to pinpoint which tRNAs are affected by 5-FU, how this influences translation and how these changes are achieved mechanistically. For this, quantification of tRNA modifications with temporal resolution is required.

### 5-FU reprograms tRNA modifications independently of transcription

We know from our previous studies in *S. cerevisiae* that chemical stressors lead to halted transcription during the acute phase of exposure. To study the transcription activity during 5-FU treatment, we designed a pulse-chase NAIL-MS experiment. For this, cells were grown in isotopically labeled medium for one week to ensure that >99% of RNA nucleosides are stable isotope labeled (30). Upon medium exchange to unlabeled medium, 5-FU and/or actinomycin D (AcmD) – a transcription inhibitor – were added to the culture and samples were drawn at different time points after experiment initiation (Figure 2A). Total tRNA was prepared by PAGE and nucleosides were quantified by LC-MS/MS. The ratio of new transcripts is calculated by dividing the abundance of unlabeled (new) by unlabeled + labeled (pre-existing) canonical nucleosides (Figure 2B). Our data shows that the new transcript ratio is identical for untreated and 5-FU treated cells, while the addition of AcmD halts transcription (40). Cells co-exposed to 5-FU and AcmD show no transcription. We thus conclude that 5-FU does not inhibit transcription of total tRNA.

**Figure 2.**
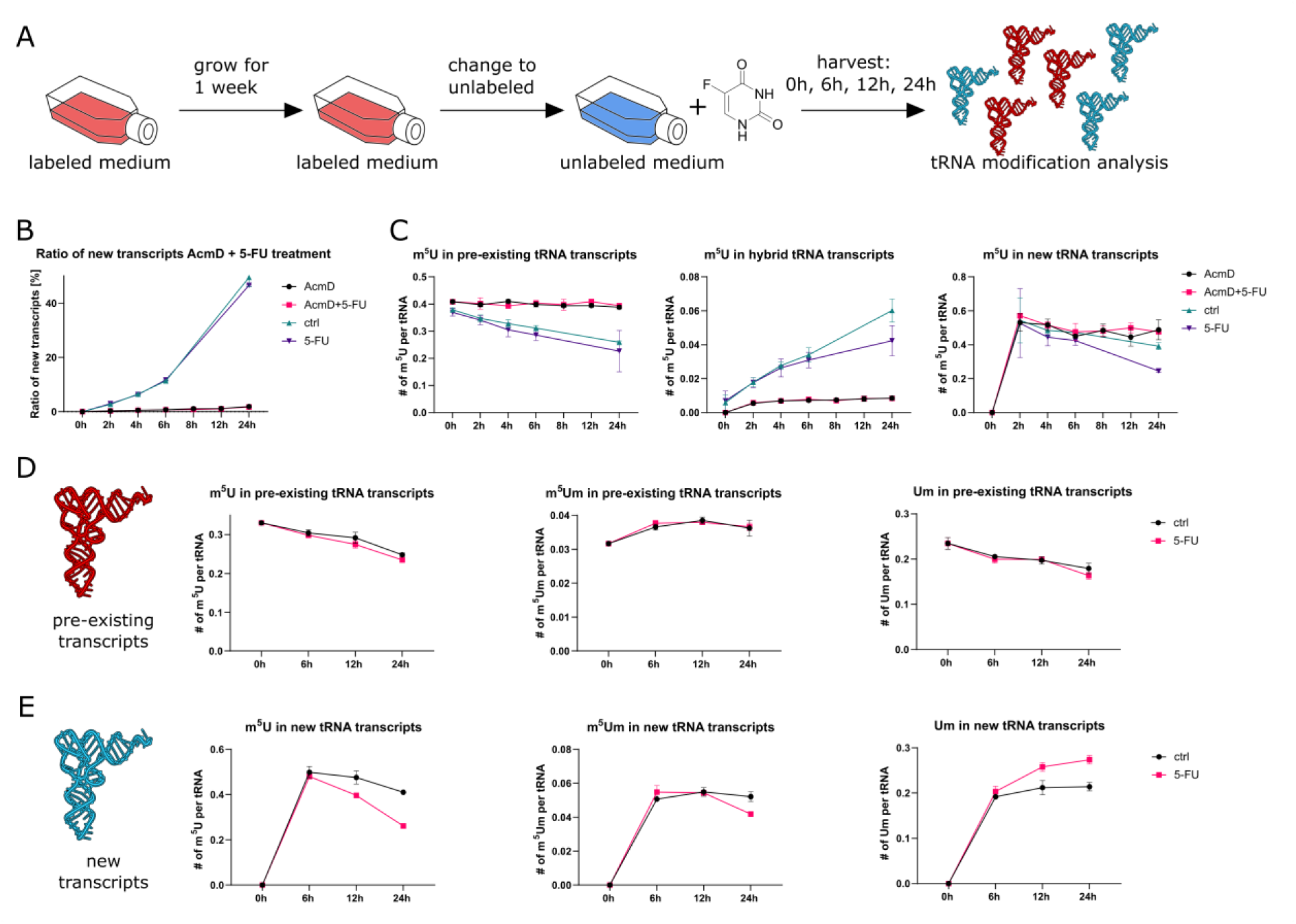
5-FU reprograms tRNA modifications independently of transcription. **A)** Concept sketch for cell culture NAIL-MS experimental design. **(B)** Ratio of new transcripts [%] under 5-FU and AcmD exposure in a time-course experiment. **(C)** Time-dependent changes of m^5^U abundance in pre-existing, new and hybrid tRNA transcripts under 5-FU and AcmD exposure. **(D)** Time-dependent changes for modifications of the m^5^U pathway (m^5^U, m^5^Um and Um) in pre-existing tRNA transcripts under 5-FU exposure. **(E)** Time dependent changes for modifications of the m^5^U pathway (m^5^U, m^5^Um and Um) in new tRNA transcripts under 5-FU exposure.

### New tRNAs exhibit altered modification abundances during 5-FU treatment

In the next step, we plotted the abundance of pre-existing and new modified nucleosides per respective pre-existing or new tRNA (referenced to the respective canonical nucleosides). For m^5^U, m^5^Um and Um, we find no difference between control and 5-FU exposed cells in tRNAs existing prior to 5-FU exposure (Figure 2C and 2D). In contrast, new tRNAs, transcribed during 5-FU exposure, show lower abundance in m^5^U and m^5^Um at later time points while Um abundance is increased (Figure 2E). In combination with our finding that it takes 6 hours for 5-FU incorporation (Figure S3), Figures 2C-E support our hypothesis for 5-methylation independent Um54 formation. In summary, the decrease in m^5^U and m^5^Um is limited to new tRNAs, as are the changes observed for other modifications in the acute phase of exposure (Figure S5). Regarding the hypermodified uridines found at position 34 of tRNA, we observe elevated levels for ncm^5^U, mcm^5^U and mcm^5^s^2^U after 24h of treatment (Figure S5).

For improved temporal resolution of NAIL-MS experiments, we included stable-isotope labeled methionine with a CD_3_-methyl group (CD_3_-methionine) in the pulse-chase set-up. With the addition of CD_3_-methionine, we can observe hybrid modifications, defined by a labeled nucleoside core structure and addition of a CD_3_-methyl group compared to a pre-existing modification which has the labeled nucleoside core but a CH_3_-methyl group. Biologically, these hybrids occur either through methylation of hybrid tRNAs, formed at the early stage of the pulse-chase experiment where both the labeled and unlabeled nucleotide pools are available to the polymerase, or through “post-methylation” of pre-existing tRNAs. To distinguish the biological processes, the RNA polymerase inhibitor AcmD is used. As shown in Figure 2C, hybrid-m^5^U is less abundant once transcription is blocked. In addition, pre-existing m^5^U remains constant as it is not diluted by hybrid tRNAs. This means that hybrid-m^5^U are not post-methylation events but a reflection of very young tRNAs, that were transcribed during early phases of 5-FU exposure. As seen in the middle panel of Figure 2C, the abundance of hybrid-m^5^U from control and treated cells is identical within the first 4 hours of 5-FU treatment and a decrease in hybrid-m^5^U abundance becomes detectable only after 6 hours of exposure. This shows that 5-FU must be incorporated into the tRNA to inhibit TRMT2A and, further, that it takes approximately 6 hours for effects to become detectable in the early hybrid-species. This explains why the abundance of m^5^U in the new transcripts (Figure 2E) is comparable at the early 2- and 4-hour time points and effects become only visible 6 hours after 5-FU exposure. Additional evidence for the 5-FU transcription-dependent effect is found in the new tRNAs taken from cells co-exposed to both 5-FU and AcmD. Here, m^5^U does not drop over time, but stays constant, which strongly argues against demethylation of m^5^U in the context of 5-FU treatment.

All in all, our pulse-chase NAIL-MS data make us confident that the drop in m^5^U (and m^5^Um) abundance is only possible due to incorporation of 5-FU into tRNA which traps TRMT2A.

### Cells reprogram their new tRNAs most pronounced during recovery from 5-FU exposure

We have found that tRNA modification abundance is only affected in new tRNAs during 5-FU treatment while pre-existing tRNAs are not affected. In Figure 1A, we have also studied the recovery period after removal of 5-FU and found that especially modifications at the wobble position (34) change during recovery. To study this observation in more detail, we performed a pulse-chase NAIL-MS experiment in which cells were grown in fully labeled medium for 7 days and continued to grow in fully labeled medium during a 24-hour exposure to 100 µM 5-FU. In the chase phase, 5-FU was removed by medium exchange into unlabeled medium and samples were drawn after 6 and 12 hours (Figure 3A). After 24 hours of 5-FU exposure, we found lower modification abundances for m^5^U and m^5^Um and higher abundances for Um in the pre-existing tRNAs (Figure 3B). For new tRNAs, we observe a continued low abundance of m^5^U which indicates that TRMT2A is still inhibited even 12 hours after removal of 5-FU. This data supports the reports of the covalent inhibition of TRMT2A through 5-FU (11,26).

**Figure 3.**
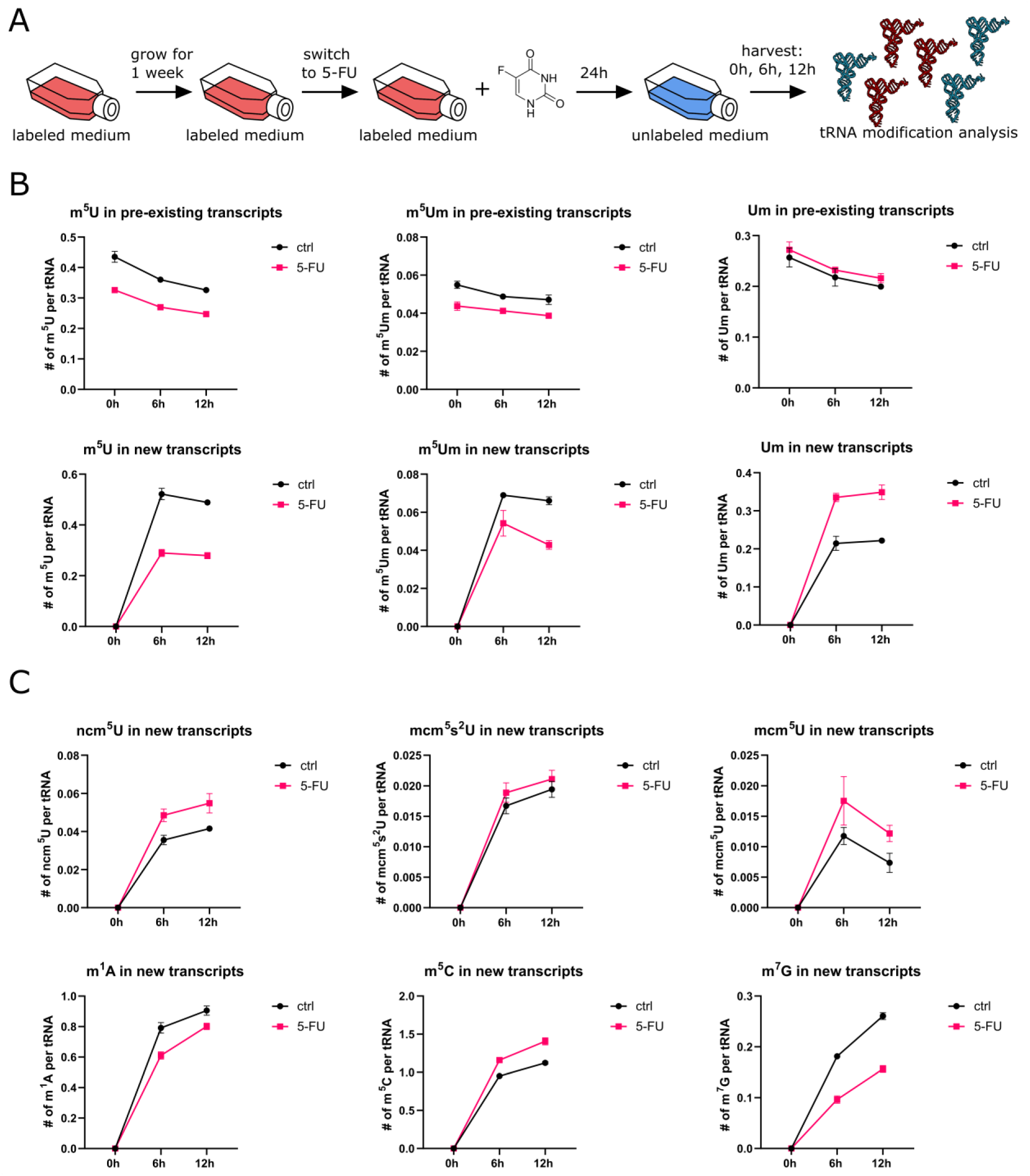
Pulse-chase NAIL-MS experiment while cells recover from a 24 hours 5-FU exposure. **(A)** Concept sketch for cell culture NAIL-MS experimental design. **(B)** NAIL-MS results for m^5^U, m^5^Um and Um in both pre-existing and new total tRNA transcripts after 24h 5-FU exposure during recovery. **(C)** NAIL-MS results for ncm^5^U, mcm^5^s^2^U, mcm^5^U, m^1^A, m^5^C and m^7^G in new total tRNA transcripts after 24h 5-FU exposure during recovery.

Next, we studied the behaviour of the hypermodified uridine derivatives, as they might be also affected by 5-FU treatment. In the pre-existing tRNAs, we observe no difference in abundance of mcm^5^s^2^U, mcm^5^U or ncm^5^U in tRNA of both untreated and treated cells (Figure S6). For mcm^5^U, a decrease is observed within the first 6 hours – which is explained by its nature as an intermediate of mcm^5^s^2^U synthesis. In this timeframe, the majority of mcm^5^U is thiolated to mcm^5^s^2^U (Figure S6). Interestingly, we find an elevated abundance of hypermodified uridines ncm^5^U and mcm^5^U in new total tRNA after 5-FU treatment. For mcm^5^s^2^U, we can find no significant increase in the new tRNA pool, although there is a trend towards higher levels of this uridine modification, too. Similar to hypermodified uridines, the abundance of other, non-uridine modifications is altered as well. The abundance of m^5^C is elevated during 5-FU recovery, while m^1^A and m^7^G are less abundant in tRNAs derived under 5-FU treatment compared to the controls (Figure 3C). For other modifications, such as m^1^G, m^22^G, m^3^C or Cm, similar changes towards up- or downregulation can be found (Figure S6). This finding contrasts the data from Figure 1A and Figure S4, where we report no or only minor changes for *e.g.* m^1^A or m^22^G. While this appears to be a contradiction in our data at first glance it, in fact, displays the power of NAIL-MS to deconvolute tRNA modification data: In the unlabeled experiment of Figure 1A, changes in new, but low-abundant tRNAs during the recovery phase are diluted by the high-abundant pre-existing tRNAs with an unaltered modification profile. Therefore, the adaptation of human cellular tRNAs as a consequence of stress exposure is only visible after the distinction of pre-existing and new tRNAs.

The fact that modifications in new tRNAs are either increased (hypermodified uridines, m^3^C, m^5^C and Cm) or decreased (m^1^A, m^22^G, m^7^G and m^1^G) can be explained by two hypotheses: (1) the stoichiometric abundance of the modification changes or (2) the abundance of isoacceptors changes during the recovery phase. To further explain the data and determine the mechanisms behind the observed tRNA modification adaptation, isolation and detection of tRNA isoacceptors are needed.

### tRNA modification reprogramming is tRNA isoacceptor-specific

From platforms such as modomics (3) or tmodbase (41), we know the modification profile of 32/47 human cytosolic tRNAs. Because hypermodified uridine modifications are particularly involved in fine-tuning the translation process, we wanted to further examine isoacceptors carrying these. mcm^5^s^2^U is reported in the tRNAs Lys^UUU^, Arg^UCU^, Gln^UUG^ and Glu^UUC^ (42), ncm^5^U in Val^UAC^, mcm^5^Um in Sec^UGA^ and mchm^5^U in Gly^UCC^ (43). From these, we chose to study tRNA^Lys^_UUU_ as a carrier of mcm^5^s^2^U and mcm^5^U as intermediate of mcm^5^s^2^U in detail. In addition, we examined the modification profile of other isoacceptors as a control (for detailed information see Figure S7, Figure S8 and Figure S9).

tRNA isoacceptors were purified using reverse-complementary, biotinylated DNA probes (Table S3) and streptavidin-coated magnetic beads. The complete modification profile is displayed in Figure S7 and it is in good agreement with literature in both chemical variety and stoichiometry. Figure 4A shows the abundance of different modifications found in tRNA^Lys^_UUU_ isolated from cells after (24h exposure + 6h recovery) 5-FU treatment. Although these cells were grown outside the NAIL context an increase in mcm^5^U is observed. This pattern mirrors the elevated levels observed in total tRNA after 24h of 5-FU exposure (Figure 1A). For the other modifications, including m^1^A, m^7^G, m^5^C and mcm^5^s^2^U, no significant changes were detected. Here, a closer inspection of tRNA^Lys^_UUU_ taken from a pulse-chase NAIL-MS experiment is needed to deconvolute the mixture of pre-existing and new tRNAs. As shown in Figure 4B, new tRNAs from recovering cells show similar abundances of mcm^5^s^2^U in tRNA^Lys^_UUU_. This indicates that the stoichiometry of mcm^5^s^2^U modification per tRNA is independent of the treatment. Detection of mcm^5^U in tRNA^Lys^_UUU_ proved challenging due to the comparably high limit of detection and low abundance (in new transcripts) of this modification. Aside from mcm^5^U and mcm^5^s^2^U, this analysis confirms the changes for m^5^U, m^5^Um, Um and Ψ we observed in total RNA (Figure S9). Consistent with our hypothesis of Um formation instead of m^5^Um, we observe elevated levels of Um upon 5-FU treatment exclusively in isoacceptors that carry m^5^Um (Figure S7). Additionally, we observed a decrease in m^7^G and m^1^A levels in tRNA^Lys^_UUU_ (Figure 4B), consistent with our recovery NAIL-MS data (Figure 3C). However, no increase in m^5^C was detected, in contrast to findings in total tRNA.

**Figure 4.**
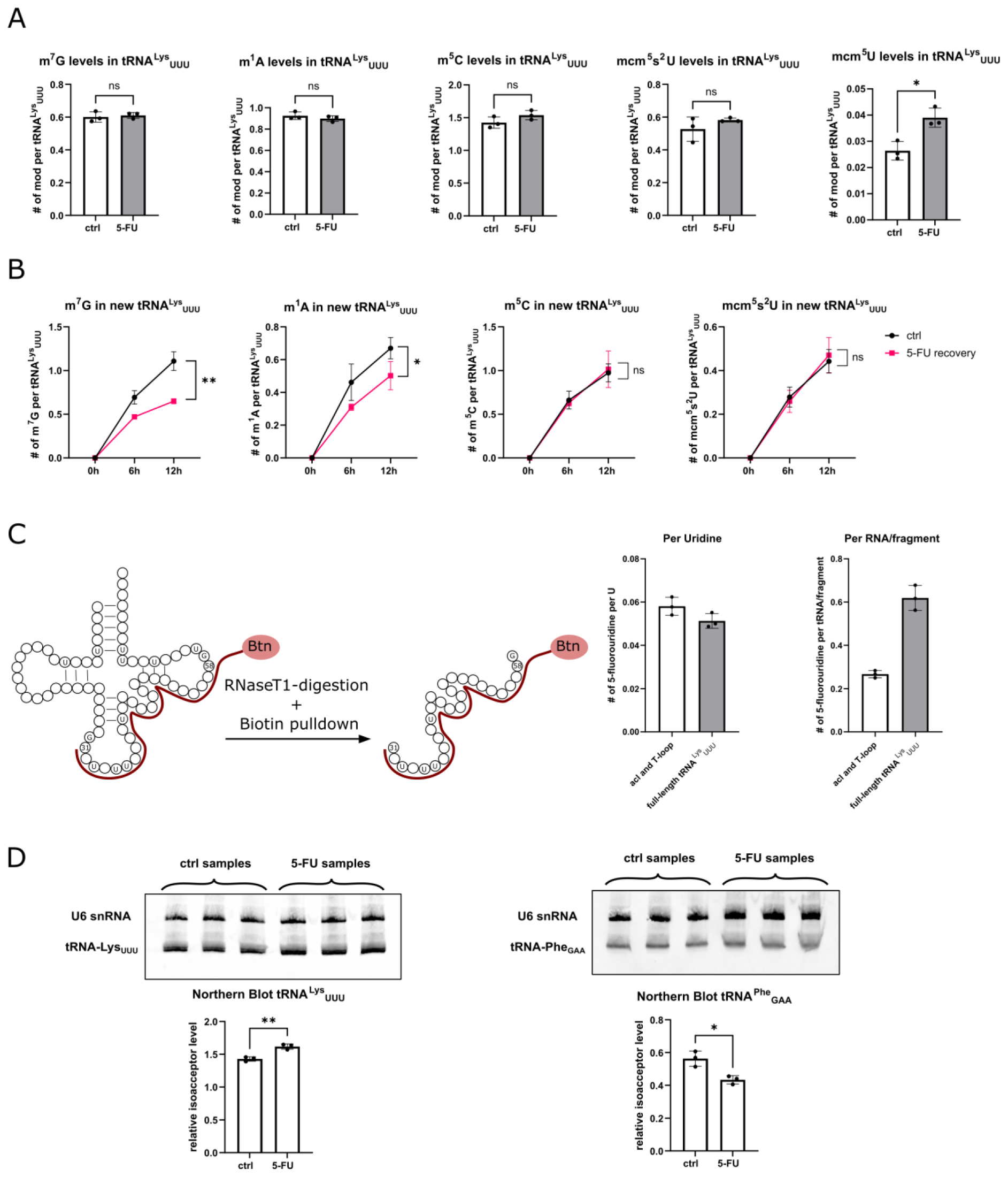
tRNA isoacceptor abundance and modification abundances in tRNA^Lys^_UUU_ after 5-FU exposure. **(A)** Modification abundances of typical tRNA^Lys^_UUU_ modifications after 24h acute 5-FU stress and 6h 5-FU recovery **(B)** Modification abundances in new tRNA^Lys^_UUU_-transcripts during recovery from 24h 5-FU exposure (6h and 12h). Data is shown for m^7^G, m^1^A, m^5^C and mcm^5^s^2^U. (Statistics exclusively for 12h timepoint: Welchs t-test, p > 0.05 (ns), p < 0.05 (*), p < 0.01 (**)). **(C)** 5-fluorouridine levels after 24h 5-FU exposure in tRNA^Lys^_UUU_ and in the anticodon-/T-loop fragment of tRNA^Lys^_UUU_ obtained after RNase T1 digestion and biotin pulldown assay **(D)** Northern Blots and Isoacceptor abundances of tRNA^Lys^_UUU_. tRNA isoacceptor abundance was evaluated using U6-snRNA as a reference.

Given the detection and quantification of 5-fluorouridine in HEK293T total tRNA, we extended our analysis to the isoacceptor level. Moreover, we examined the amount of 5-fluorouridine in a fragment of tRNA^Lys^_UUU_, carrying sections of the anticodon- and the T-loop. An RNase T1 digest was performed between oligonucleotide hybridization and isolation steps, yielding a fragment spanning from position 30 to 59 of the respective tRNA (Table S3). As shown in Figure 4C, 5-fluorouridine is present in both full-length tRNA and the isolated fragment. The fragment contained less 5-fluorouridine (0.25x per fragment) compared to the full-length tRNA (0.62x per tRNA^Lys^), which is due to its lower length. By normalization to the number of Us in the RNA, we find a 6% chance of 5-FU incorporation for each U position in both the full-length and the fragment. In addition to tRNA^Lys^_UUU_, we conducted analysis on tRNA^Asn^_QUU_ and tRNA^Phe^_GAA_ and could detect and quantify 5-fluorouridine in these isoacceptors, too (Figure S15).

To further explore the effect of 5-FU on isoacceptors, we studied isoacceptor abundances using Northern Blotting (NB). Typically, 5S-rRNA serves as a reference for NB analysis; however, we observed that 5-FU alters ribosomal RNA levels during tRNA isolation by SEC (also see next section), leading us to use U6-snRNA instead of 5S-rRNA as a reference. Figure 4D shows a modest increase in the relative abundance of tRNA^Lys^_UUU_. To confirm that this elevation was not influenced by changes in the reference RNA, we also analyzed the relative abundance of tRNA^Phe^_GAA_, which showed a slight decrease in abundance. Northern Blotting of tRNA^Asn^_QUU_ and tRNA^Glu^_UUC_ (Figure S8) revealed similar trends, with elevated tRNA^Asn^-levels and reduced tRNA^Glu^-levels. However, our analysis of the latter two isoacceptors lacked statistical significance and should be interpreted with care.

Even though these isoacceptor level changes are modest, they point to the complex impact of 5-FU on specific tRNAs. Modification changes in new tRNA transcripts during recovery from 5-FU may parallel modification changes in the specific isoacceptors. Conversely, the relative abundance of individual isoacceptors seems to fluctuate, possibly due to differential decay of some isoacceptors, as already described for fluorinated pyrimidines or 5-FU in yeast and HeLa cells, respectively (44,45).

### 5-FU changes abundance of ribosomal RNAs and their modification profile

As we investigated the abundance of tRNA isoacceptors, we became curious whether the levels of different rRNA subtypes might also be affected. First, we compared size-exclusion chromatography (SEC) chromatograms of total RNA from cells treated with 5-FU for 24 hours to those from untreated cells. Despite injecting equal amounts of total RNA, the signal intensity for the 28S and 18S rRNA double-peak was reduced in 5-FU treated cells, as shown in Figure 5A. In contrast, tRNA signal intensity remained unchanged or even slightly increased.

**Figure 5.**
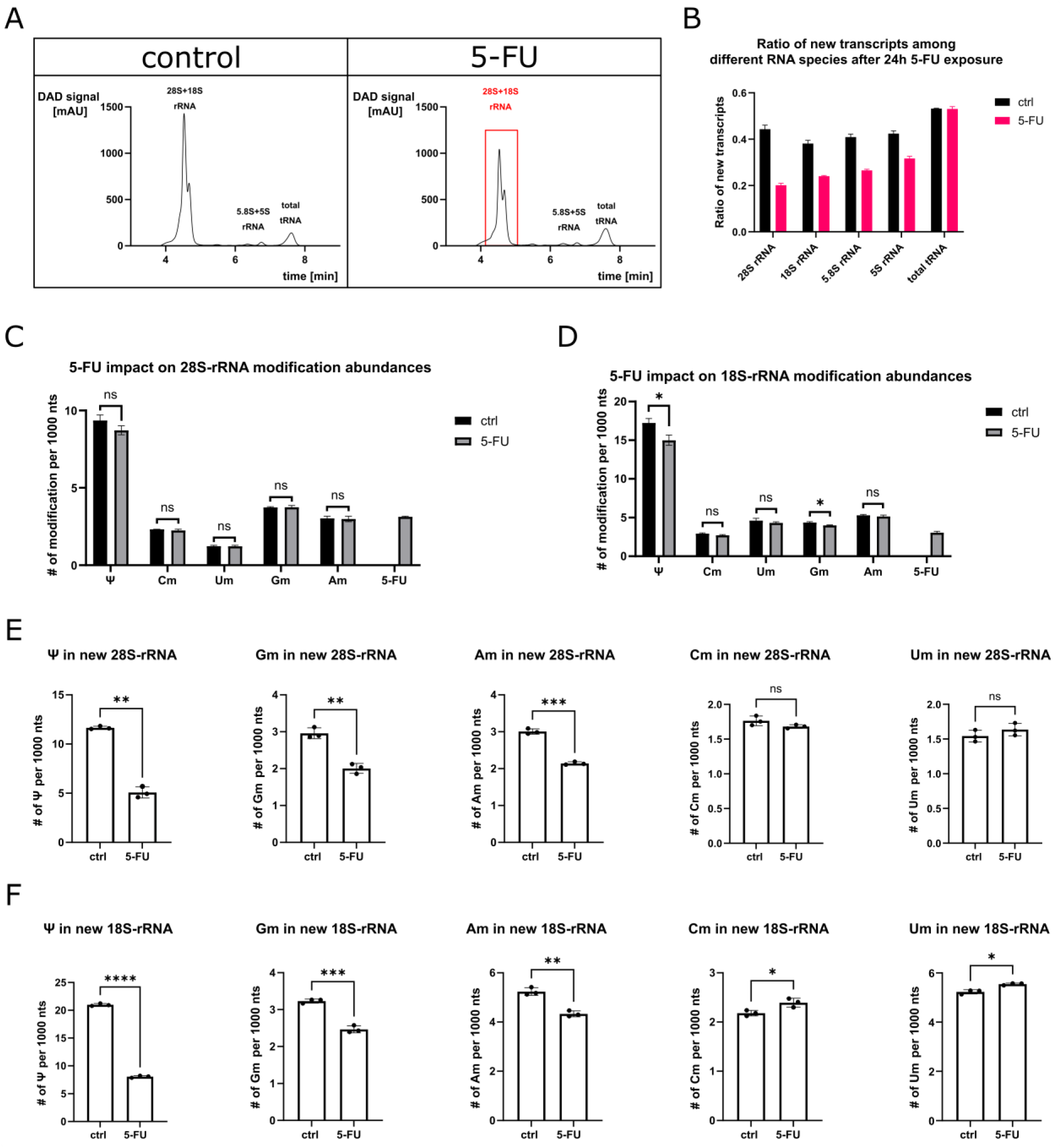
5-FU effects on RNA abundance and rRNA modification profile. **(A)** UV chromatograms (λ=254 nm) of untreated and 5-FU treated total HEK RNA obtained by SEC. **(B)** Ratio of new transcripts for different RNA species after 24h 5-FU exposure calculated via NAIL-MS. **(C)** 5-FU impact on 28S-rRNA modification abundances after 24h 5-FU exposure. **(D)** 5-FU impact on 18S-rRNA modification abundances after 24h 5-FU exposure. **(E)** Modification abundances in new 28S-rRNA transcripts after 24h 5-FU exposure obtained by NAIL-MS. **(F)** Modification abundances in new 18S-rRNA transcripts after 24h 5-FU exposure obtained by NAIL-MS. (Statistics: Welchs t-test, p > 0.05 (ns), p < 0.05 (*), p < 0.01 (**), p < 0.001 (***), p < 0.0001 (****)).

Given the decrease in rRNA subunits, we examined the ratio of newly synthesized rRNA transcripts compared to the respective total rRNA amount in a NAIL-MS experiment (sum of new and pre-existing transcripts). After 24 hours of 5-FU exposure, the proportion of all new rRNA transcripts dropped by approximately 30–50% (Figure 5B) compared to untreated cells, while the ratio of new tRNA transcripts was unaltered (Figure 2B). This observation agrees with the literature. Studies across different organisms have shown that 5-FU primarily affects rRNA maturation, leaving the transcription of precursor rRNA species (47S/45S-rRNA) unaffected, while rRNA processing into 28S-, 18S- and 5.8S-rRNA is impaired (28,29,46).

Intrigued by these findings, we explored whether 5-FU also alters modification patterns in rRNA. Following 24 hours of 5-FU treatment, we isolated 28S- and 18S-rRNA via SEC and conducted LC-MS/MS analysis. These two subtypes were selected due to their extensive and diverse modification profile compared to 5.8S- and 5S-rRNA. For 28S rRNA modifications, no significant changes were observed (Figure 5C). Notably, our analysis showed a significant reduction in Ψ and Gm levels in 18S-rRNA after the treatment (Figure 5D). Other common modifications in 28S- and 18S-rRNA remained unaffected (Figure S10). However, 5-fluorouridine was successfully detected in both subunits, leading us to the assumption that processing is partially maintained.

For total tRNA, we found that only new tRNAs have altered modification abundances under 5-FU treatment. To test whether this is also true for rRNA, we purified the 28S- and 18S-rRNA from the NAIL-MS experiment in Figure 5B to differentiate between pre-existing and newly synthesized transcripts. In new 28S-rRNA transcripts, levels of Ψ, Gm and Am are lower after 24 hours of 5-FU exposure (Figure 5E), a pattern also observed in 18S-rRNA (Figure 5F). Interestingly, additional modifications in 18S-rRNA, such as Cm, ac^4^C and Um, showed slight increases, while m^6,6^A levels decreased (Figure 5F and Figure S11).

To determine whether these modification changes were due to RNA degradation, we assessed RNA integrity via chip gel electrophoresis. Based on established protocols (47), we further examined if marker modifications of 28S-rRNA, like m^3^U, were present in the respective 18S-rRNA fraction. The RNA integrity number (RIN) values (Figure S12) indicated a good RNA quality. Additionally, no characteristic degradation-associated modifications were found within the 18S-rRNA pool. Thus, the observed changes in modification levels are not due to degradation but are likely a direct result of 5-FU treatment.

### Proteins are differentially expressed during and after 5-FU exposure

We found distinct changes in translation-modulating RNA modifications and RNA abundances in our studies. These changes might influence the translation of proteins and therefore we performed shotgun proteomics of cells grown for 24 hours in the presence of 5-FU and cells stressed with 5-FU after a 6h recovery period after 5-FU removal. We identified approximately 1600 protein groups, respectively (Figure 6A and Table S4). Under acute 5-FU treatment, we found 21 proteins to be differentially expressed in 5-FU cells compared to the untreated controls. While 7 proteins were significantly more abundant, 14 proteins appear to be significantly less abundant. The lower-abundant proteins mainly contain 60S and 40S ribosomal proteins, consistent with existing literature (48).

**Figure 6.**
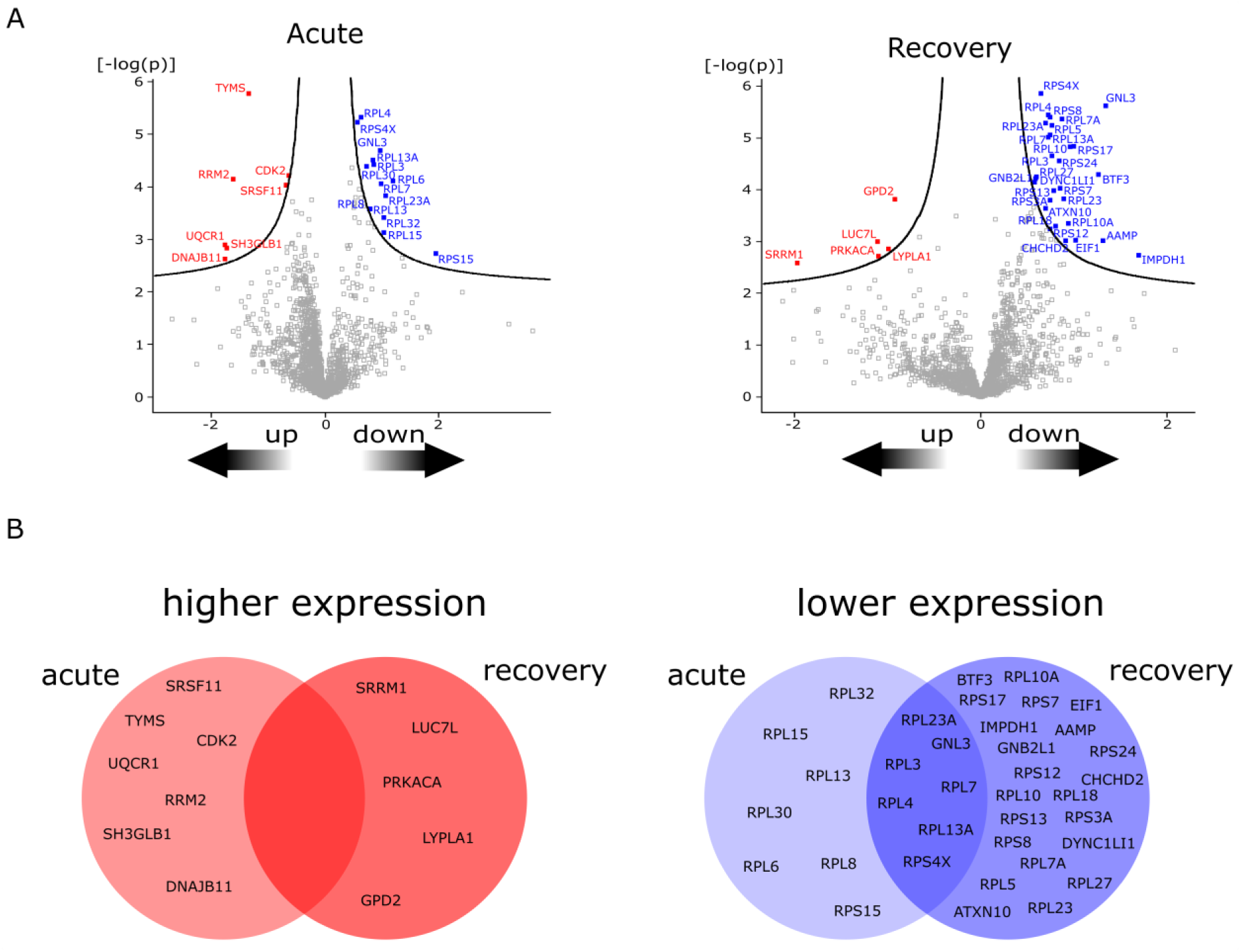
Protein expression during and after 5-FU stress. **(A)** Volcano plots for proteins being up- and downregulated after 24h 5-FU exposure (acute - left) and 24h 5-FU exposure with 6h wash-out in fresh medium (recovery - right). **(B)** Venn diagram showing similar changes in protein expression between 24h acute exposure (left) and after 24h exposure following 6h recovery (right).

After 6 hours of recovery, we found 34 proteins that significantly differ in abundance compared to the untreated control. Now, only 5 proteins are more abundant, while 29 proteins, again mainly 60S/40S ribosomal proteins, are less abundant. A comparison of up- and downregulated proteins identified during acute and recovery conditions revealed that 7 proteins are lower abundant in both conditions while no proteins in the upregulated group were shared (Figure 6B). Our analyses detected 28 proteins involved in tRNA maturation and modification (e.g. ELP3, NSUN2, TRMT1 or diverse tRNA ligases, Table S1 and S2)), but no statistically relevant differences. This indicates that the changes in tRNA modifications such as mcm^5^(s^2^)U (writer: ELP3), m^5^C (writer: NSUN2), m^7^G (writer: METTL1) or m^22^G (writer: TRMT1) are not caused by changed abundance of the respective tRNA-writers.

Previous studies have highlighted the crucial role of 5-FU in translational reprogramming, particularly through its association with fluorinated ribosomes (49). Given the reprogramming of different RNA modifications in both tRNA and rRNA, we aimed to further explore the link between the modification level and translational reprogramming. One common approach for investigating differential translational activity is through gene specific codon usage analysis (GSCU) and tRNA isoacceptor usage (IAU) analysis of up- and downregulated proteins (14). For this purpose, we compared the GSCU and IAU of the upregulated proteins to all human protein-coding genes and especially during recovery, we found statistically significant differences in both GSCU and IAU (Figure S13 and S14). But given the fact that our proteomics analysis revealed only 5 upregulated proteins, the biological significance of the GSCU and IAU analysis might be modest and should be interpreted with great care.

## DISCUSSION

In recent decades, numerous studies have explored the effects of 5-fluorouracil (5-FU) on RNA biology across various stages. It has long been established, spanning over 40 years, that 5-substituted uridine modifications are particularly responsive to 5-FU treatment (22). This reduced abundance arises from two primary mechanisms: the incorporation of 5-fluorouridine instead of uridine into RNA and the inhibition of writer enzymes such as TRMT2A and pseudouridine synthases (26,27).

In our investigation, we adopted the current state-of-the-art approach to examine tRNA modification levels in HEK293T cells, utilizing absolute quantification through LC-MS/MS. These findings confirm the loss of m^5^U and Ψ. Furthermore, we found significant alterations in other modification abundances within the m^5^U pathway. Specifically, we observed changes in m^5^Um and Um, indicating a shift from m^5^U/m^5^Um to Um formation (Figures 1A and 1B). This is most likely due to the inhibition of the m^5^U writer TRMT2A, while the writer for m^5^Um (which remains unknown) appears to be unaffected by 5-FU treatment. Other modifications, *e.g.* m^1^A or m^2,2^G, show only modest to no changes within the total tRNA pool (Figure 1B). Other than that, we showed that 5-FU is incorporated into tRNA which is in accordance with hypotheses in the literature (50,51). In addition, we found 5-fluorouridine to be detectable by LC-MS/MS after approximately 6 hours of incubation (Figure S3).

To validate our findings for altered uridine modification abundances, we performed NAIL-MS which offers insights into the dynamics of RNA modification in both newly synthesized and existing tRNA transcripts. Notably, our results demonstrate that only new tRNA transcripts are affected by 5-FU treatment, and modification reprogramming occurs independently of transcription (Figure 2B). For modifications within the m^5^U-pathway, we find a significant shift from m^5^U and m^5^Um formation towards Um in new tRNA transcripts (Figure 2E). We hypothesize that 5-FU leads to Um accumulation specifically in isoacceptors carrying m^5^Um. According to this hypothesis, only the m^5^U-writer TRMT2A would be affected by 5-FU (26), while the responsible writer for m^5^Um seems unaffected by 5-FU exposure and carries out further modification of U54 to Um. However, the writer responsible for m^5^Um remains unidentified, leaving the proposed mechanism speculative at this stage. As future research may uncover the m^5^Um writer, additional experimentation could provide further insights into this mechanism.

To further dissect the impact of 5-FU on m^5^U, we advanced our NAIL-MS experiments using D_3_-methionine and actinomycin D to investigate the effects once transcription is blocked (Figure 2C). In the pre-existing transcripts, we observe a constant decrease of m^5^U abundance over time in the untreated control cells. Under AcmD treatment, this decrease is no longer observed which leads to our conclusion that the decrease is transcription-dependent and most likely caused by hybrid transcripts that contain both labeled and unlabeled nucleosides. From the hybrid tRNAs that emerge early on after medium exchange, we can see that m^5^U forms equally fast in control and 5-FU treated cells in the first 2 hours after 5-FU exposure. But then, the inhibition of TRMT2A through 5-FU becomes apparent and the m^5^U abundance rises less quickly in the 5-FU treated cells compared to the control. The same picture is seen in new transcripts. At the early 2-hour time point the abundance of m^5^U is identical but drops rapidly in the 5-FU treated cells due to the dilution of the m^5^U signal caused by ongoing transcription.

Further supporting evidence from our 5-FU recovery experiments underscores the altered dynamics of the m^5^U pathway, as decreased levels for m^5^U and m^5^Um were observed in the pre-existing transcripts alongside new transcripts (Figure 3B). In contrast, Um levels were also elevated in both transcript populations, but higher levels in pre-existing transcripts lacked statistical significance. This could be caused by a dilution effect, as transcripts influenced by 5-FU treatment (24h incubation) mix with those present before treatment.

More intriguingly, our recovery NAIL-MS analysis revealed unexpected alterations for modifications like m^7^G, m^1^A, m^5^C or wobble uridine modifications in new tRNA transcripts (Figure 3C). A prior study (45) demonstrated a link between the modifying enzymes NSUN2 and METTL1 with 5-FU sensitivity in HeLa cells, showing that the knockout of these enzymes, combined with 5-FU treatment, leads to rapid tRNA decay (RTD). A similar phenomenon was recently identified in yeast under additional heat stress (44), though in the human cell culture study no correlation between heat stress and RTD was found.

As NSUN2 and METTL1 are responsible for m^5^C and m^7^G placement in tRNA, an interconnection between 5-FU treatment and these modifications in tRNA is plausible. In our study, m^7^G levels were particularly lower after 24h of treatment during recovery, whereas m^5^C levels were elevated in the total tRNA pool. This elevation of m^5^C might represent a compensation mechanism to prevent RTD. To test this hypothesis in the future, further NAIL-MS experiments in the respective knock-out cells might be useful.

To explore the modification changes more specifically, we conducted isoacceptor level experiments focusing on tRNA^Lys^_UUU_. In experiments without labeling strategy, the expected effects on m^5^U, m^5^Um and Um were observed after 24h of 5-FU treatment (Figure S7). Interestingly, the Um increase was only seen in isoacceptors carrying m^5^Um, further supporting our hypothesis of Um formation independent of 5-uridine methylation. Other modifications in tRNA^Lys^_UUU_ show no to minor changes, e.g. mcm^5^U with a modest increase after 24h of 5-FU exposure (Figure 4A). It has been demonstrated that Trm9-catalyzed modifications, such as mcm^5^U, link translation to the DNA damage response (52,53). Since 5-FU induces DNA damage (24,54), we hypothesize that the increase in mcm^5^U might be a cellular stress response in human cells similar to the observations in yeast. However, the change is modest, and not reflected in the more abundant final product mcm^5^s^2^U.

We proceeded to isolate tRNA^Lys^_UUU_ from our NAIL-MS studies. During recovery, after 24h of 5-FU exposure, we found decreased levels for m^1^A and m^7^G, which is consistent with our results from total tRNA (Figure 4B). Notably, the reduction in m^1^A is intriguing, as m^5^U, m^5^Um and Um also showed altered abundances in another study with TRMT6 knock-out cells lacking m^1^A58 (55). A quite similar study further analyzed these modifications in the T-loop of tRNA^iMet^ in yeast – offering valuable insight into the cross-talk between m^1^A, Ψ and m^5^U (56). As can be seen within our results, all of these three modifications are underrepresented in new tRNA^Lys^_UUU_ which might be connected to a similar crosstalk of tRNA modifications in human cells.

We detected 5-FU incorporation as 5-fluorouridine in total tRNA, and subjected tRNA^Lys^_UUU_ to LC-MS to quantify the amount of 5-FU in this specific isoacceptor. In summary, we found that each position has a probability of ∼6% to carry 5-FU instead of U. This indicates a random incorporation of 5-FU into the RNA and a probability of 6% that U54 will be 5-FU and thus inhibit TRMT2A as hypothesized by others (9,26).

To investigate the effect on isoacceptor abundance, we performed Northern Blotting on tRNA^Lys^_UUU_ and tRNA^Phe^_GAA_. Using U6-snRNA as a housekeeper, we found elevated levels of the lysine-isoacceptor and reduced levels of the phenylalanine-isoacceptor (Figure 4D).

Similar trends were observed for other isoacceptors (Figure S8), though these changes were not statistically significant. On one hand, overexpression of isoacceptors following 5-FU treatment has been previously reported in *S. pombe* (57) - on the other hand, a rapid tRNA decay (RTD) for tRNA^Val^_AAC_ was observed under 5-FU stress when the modifying enzymes (METTL1 and NSUN2) were knocked out in *S. cerevisiae* (44) and HeLa cells (45), respectively. Although isoacceptor abundance changes in our experiments were minor, we hypothesize that 5-FU treatment alters the overall level of individual isoacceptors within the total tRNA pool, potentially driven by tRNA decay induced by tRNA hypomodification. This lower abundance of some tRNA isoacceptors will lead to a relative over-abundance of other tRNAs, although these might, in fact, stay unaltered. DORQ-seq, a recently published technology for tRNA quantification, might be a future solution to decipher these effects in detail (58).

Given the general changes in RNA abundance, we also examined rRNA levels. Literature suggests that rRNA maturation and processing are impaired under 5-FU treatment (28,29), and our Northern Blot experiments revealed reduced 5S-rRNA levels. Consistent with our size-exclusion chromatography data showing reduced rRNA signal intensities (Figure 5A), all new rRNA transcripts displayed decreased abundances after 24h of 5-FU exposure in our NAIL-MS studies (Figure 5B). These experiments also uncovered distinct changes in rRNA modification patterns in new rRNAs, including reduced levels of pseudouridine, as well as alterations in the methylation landscape. Specifically, Am and Gm levels were lower in both 28S and 18S rRNA, while Cm and Um levels were slightly elevated exclusively in 18S rRNA (Figures 5E and 5F). Importantly, these changes are only observed in the NAIL-MS context and not in the unlabeled experiments (Figure 5C and D). This is due to the high abundance of pre-existing rRNAs, that overshadow the changes that happen on the new rRNAs. From our data, we hypothesize that both 5-FU incorporation and reprogrammed rRNA methylations contribute to impaired rRNA processing. Although different studies claimed severe inhibition of rRNA maturation, we demonstrate the presence of 5-FU in processed 28S- and 18S-rRNA, indicating that rRNA processing is at least partially maintained. This observation aligns with previous reports (34,49). Another study found that the functionality of the processing RNP complex, including U3-snoRNA, remained unaffected by 5-FU (28). This also suggests that processing defects may result from 5-FU incorporation and impaired modifications such as 2’-O-methylation or pseudouridylation. Nevertheless, with only MS-based rRNA modification profiling, identifying the precise mechanism behind this remains challenging. A potential solution may involve spatially resolved analysis, where distinct cellular compartments (e.g. nucleolus, nucleus, cytosol) are individually examined by different approaches. In addition to pre-rRNA processing, rRNA modifications are crucial for translational fidelity and efficiency (59–61). While our proteomics study revealed only few differentially expressed proteins, we observed a general decrease in many ribosomal proteins (Figures 6A and 6B). Altered ribosomal protein abundance has also been reported in previous studies (48). Changes in ribosome function following 5-FU treatment were further linked to altered translation (49). In that study, polysome profiling revealed a translational reprogramming through fluorinated ribosomes. When we compared the differentially expressed proteins from our experiments to these ribosome profiling results (Table S5), we found similar trends in ribosomal protein changes.

Interestingly, the same study highlights translational upregulation of cell survival associated mRNAs. Specifically, the survival protein IGF-1R has been shown to promote 5-FU drug tolerance, which can be linked to altered translation through the fluorinated ribosome. If ribosome incorporation of 5-FU could be hindered, this might enhance the overall efficacy of the drug in affecting cancer cell fate. Other studies have similarly demonstrated that ribosome biogenesis is critical for cancer drug tolerance. For instance, in p53-inactivated cancer cells, ribosome biogenesis is hyperactivated, as p53 suppresses the expression of rRNA methyltransferase fibrillarin (62). If p53 activation remains intact, this could explain our observation of reduced rRNA methylation following 5-FU treatment. Furthermore, studies have pointed out the link between ribosomal stress and p53, suggesting a mechanism for p53 activation mediated by 5S-rRNA, RPL5 and RPL11 (63,64). Another group has shown that RPL27A protein levels, which are also reduced due to 5-FU treatment, can be restored with the addition of martynoside (65). In our proteomics data, we observed a reduction in RPL5 levels during recovery from 5-FU treatment. Additionally, our analysis revealed a decreased abundance of 5S-rRNA. Thus, our findings align well with existing data, and our NAIL-MS analysis provides a modification-centric insight into some of the mechanistic patterns affecting 28S- and 18S-rRNA.

In summary, we explored the dynamic alterations within the epitranscriptome caused by 5-FU. While it has long been established that 5-FU impacts RNA at various levels, our findings highlight the detailed view that NAIL-MS can provide to RNA modification reprogramming. This technique holds potential for addressing even more complex questions in the future, such as how impaired RNA modification influences drug resistance in different cells. Given the mechanistic relevance of our observations, this line of research may contribute to understanding and overcoming other, drug-related side-effects in the coming years.

## DATA AVAILABILITY

*The proteomics data underlying this article were deposited in the ProteomeXchange Consortium PRIDE* [https://www.ebi.ac.uk/pride; *dataset identifier: PXD051672 Project DOI: 10.6019/PXD051672]; Nucleoside LC-MS data are available in the article and in its online supplementary material*.

## SUPPLEMENTARY DATA

Supplementary Data are available at NAR online.

## AUTHOR CONTRIBUTIONS

Maximilian Berg: Conceptualization, Formal analysis, Methodology, Validation, Writing— original draft. Chengkang Li: Formal analysis, Methodology. Stefanie Kaiser: Conceptualization, Formal analysis, Visualization, Writing—original draft, review & editing.

## ACKNOWLEDGEMENTS

The authors want to thank the MS Service Facility at Campus Riedberg, especially Leona Rusling.

## FUNDING

This work was supported by the Deutsche Forschungsgemeinschaft [325871075-SFB 1309 and 259130777-SFB 1177 to S.K.]. Funding for open access charge: Horizon 2020 Program (325871075-SFB 1309).

## CONFLICT OF INTEREST

There are no conflicts to declare.

## Notes

### Competing Interest Statement

The authors have declared no competing interest.

### Summary of Updates

- Figure 4A: The bar graph for mcm5s2U levels in tRNA-Lys was declared "m7G levels in tRNALysUUU". This was corrected to "mcm5s2U levels in tRNALysUUU" - headline of section "proteins decoded by 5-FU reprogrammed tRNA isoacceptors are differentially expressed during and after 5-FU exposure" was changed to "proteins are differentially expressed during and after 5-FU exposure"

