## Supplementary file for "Temporal placement of RNA modifications under 5-fluorouracil treatment in human cell culture"

### Supplementary information

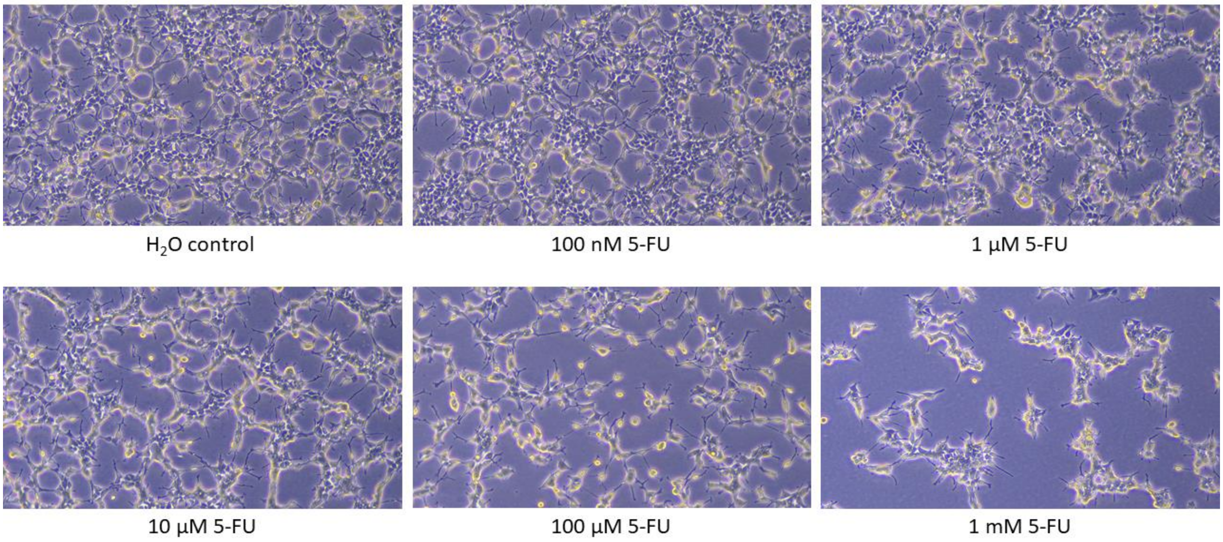


Fig. S1. HEK293T cells being cultivated for 24h with 5-FU using varying concentrations.

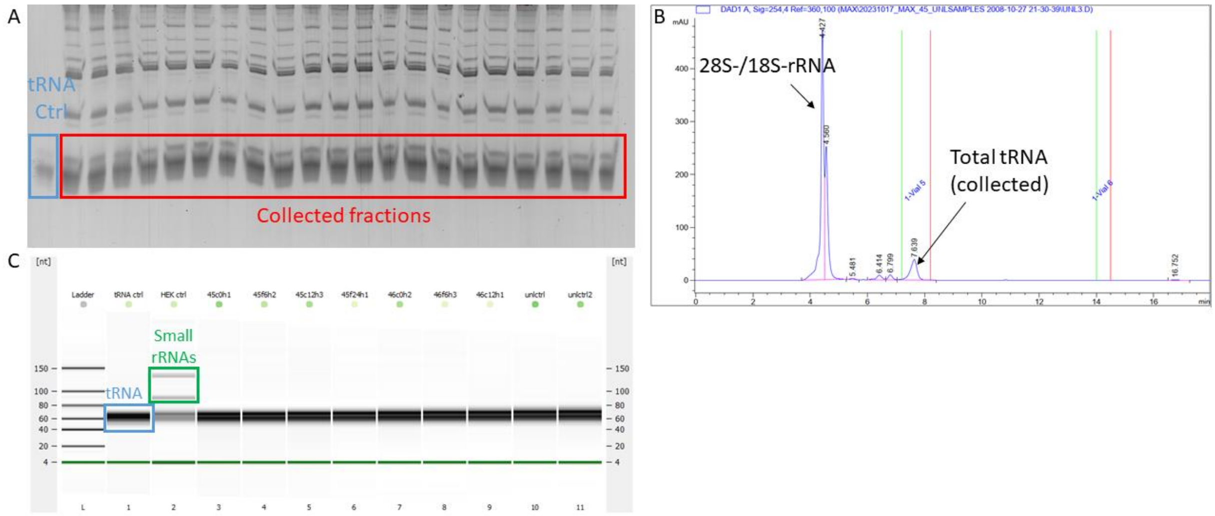


Fig. S2. Exemplary pictures for tRNA purification methods and the respective quality control (QC) (Bioanalyzer). **A** Gel picture; tRNA purification via 10% TBE-urea-PAGE, visualized using GelRed stain **B** SEC chromatogram; tRNA purification using 300 Å column **C** Agilent Bioanalyzer small chip as QC for tRNA purification.

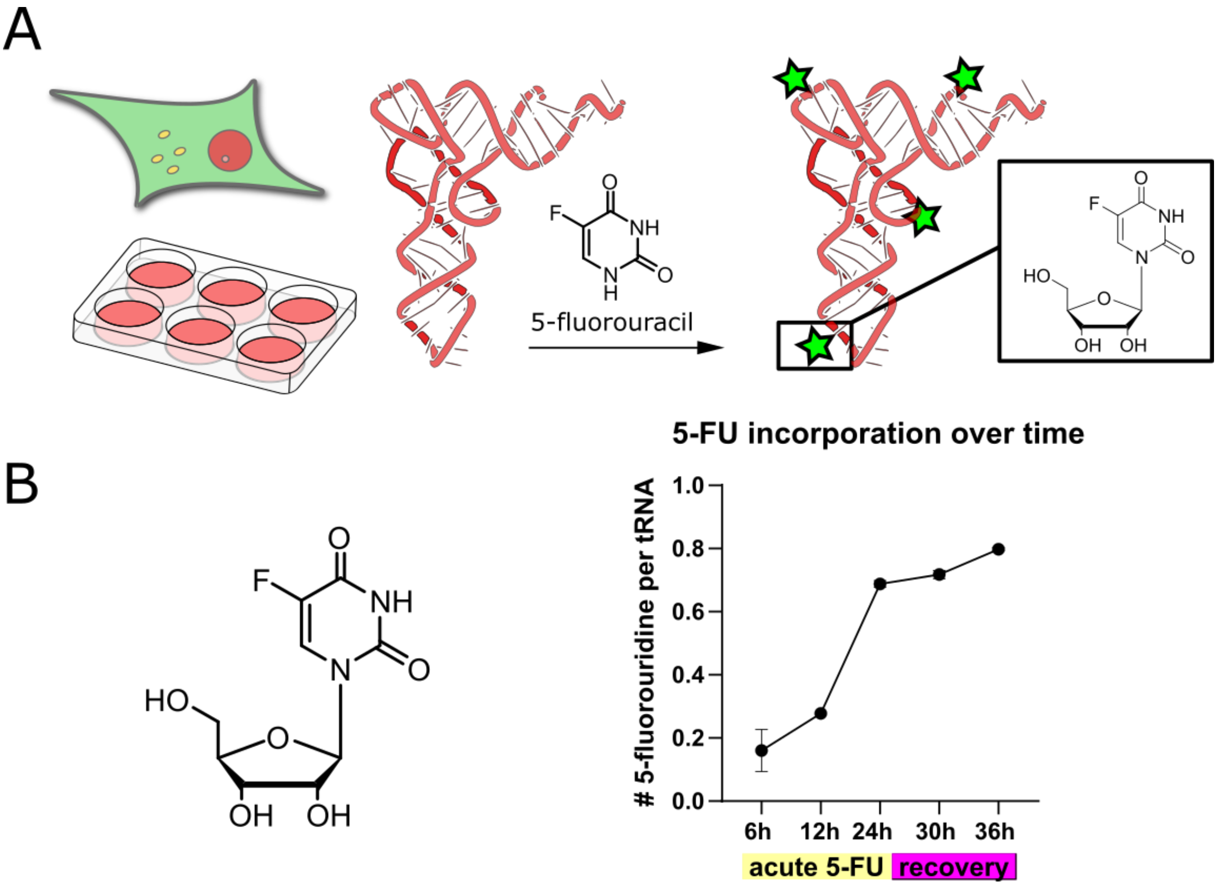


Fig. S3. Incorporation of 5-FU as 5-fluorouridine in HEK293T total tRNA. **(A)** Concept sketch for the incorporation of 5-FU in HEK293T tRNA **(B)** Number of 5-fluorouridine per HEK293T tRNA. Cells were incubated with 100 µM 5-FU and harvested after 6h, 12h and 24h, respectively. In addition, cells were pre-incubated for 24h with 100 µM 5-FU, then supplied with fresh medium and harvested after 6h and 12h recovery, respectively. 5-FU is detected as 5-fluorouridine by nucleoside-LC-MS.

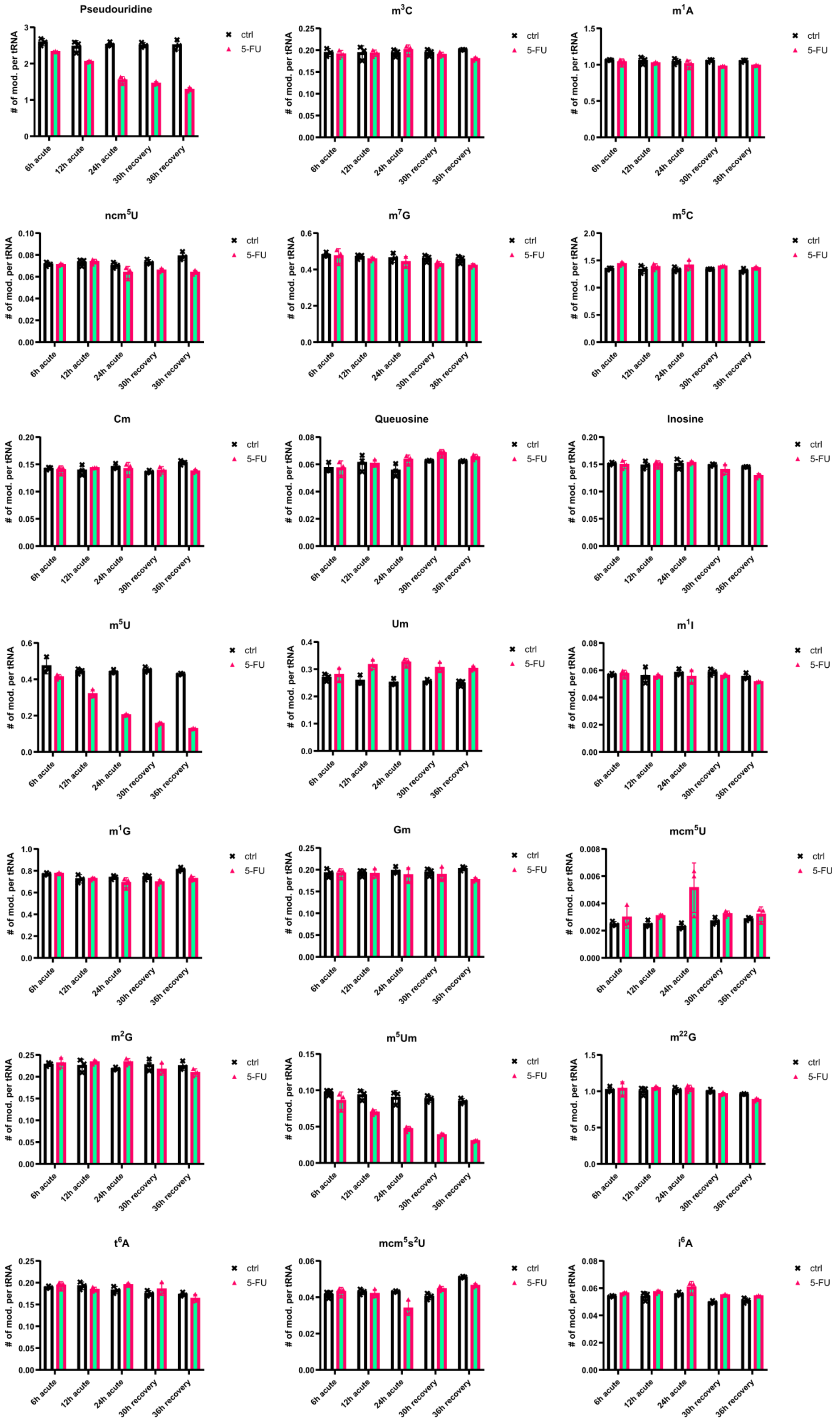


Figure S4: Modification profile of HEK293T tRNA during and after 5-FU exposure at different timepoints (6h, 12h, 24h and 24h with 6h and 12h recovery, respectively). Number of modification per tRNA was then calculated using the molar amount of modified nucleosides divided by the molar amount of tRNA. The molar amount of tRNA was calculated using the canonical nucleosides.

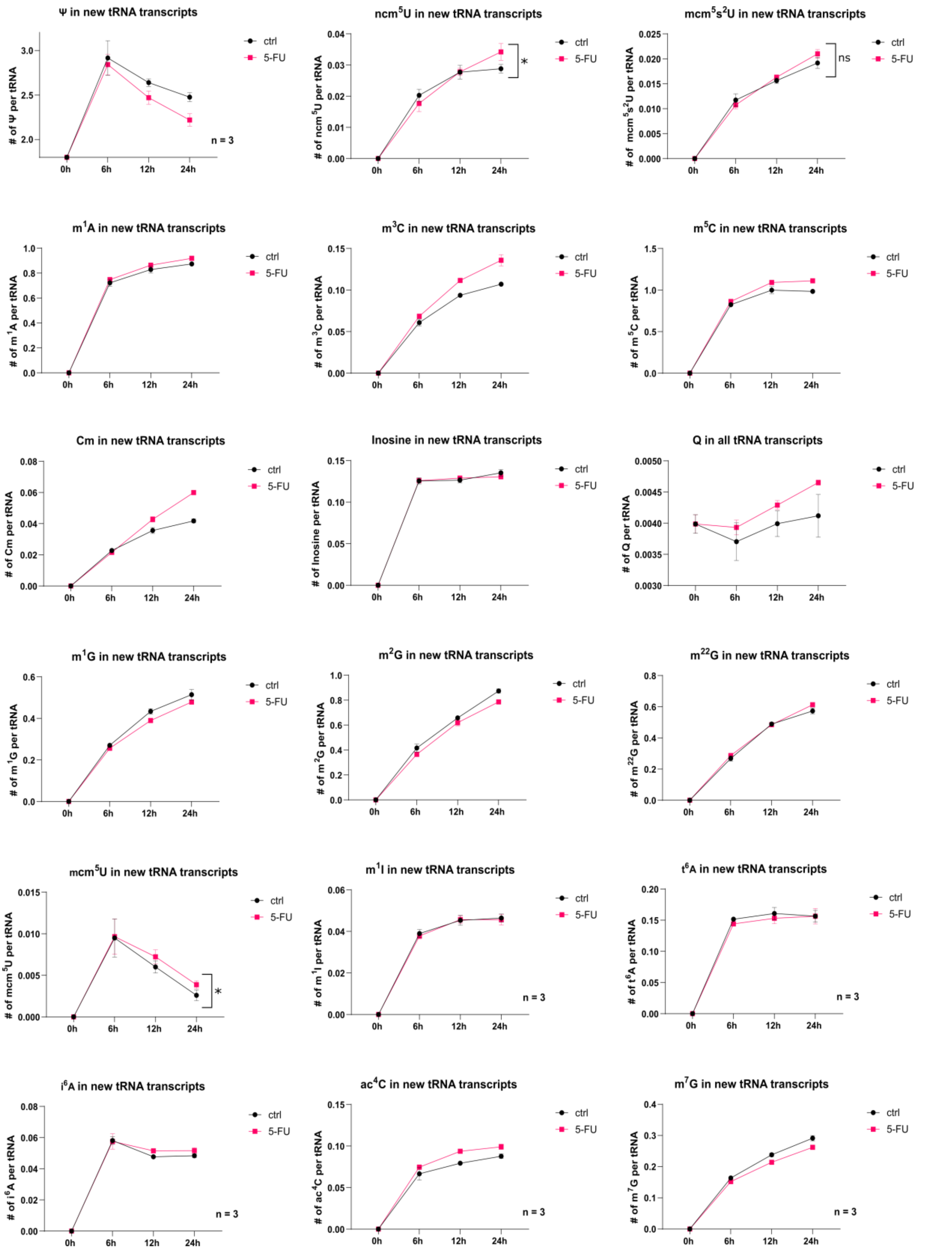


Figure S5.1: Modification profile of HEK293T new tRNA transcripts during 5-FU exposure. Cells were harvested after 6h, 12h and 24h, respectively. Modifications in new tRNA transcripts were normalized using the molar amount of unlabeled, modified nucleoside divided by the molar amount of unlabeled tRNA. Unlabeled tRNA molar amount was calculated using the unlabeled, canonical nucleosides. Note: For ncm$^5$U, mcm$^5$U and mcm$^5$s$^2$U, the statistical significance is displayed for the 24h timepoint ($P > 0.05$ is considered significant).

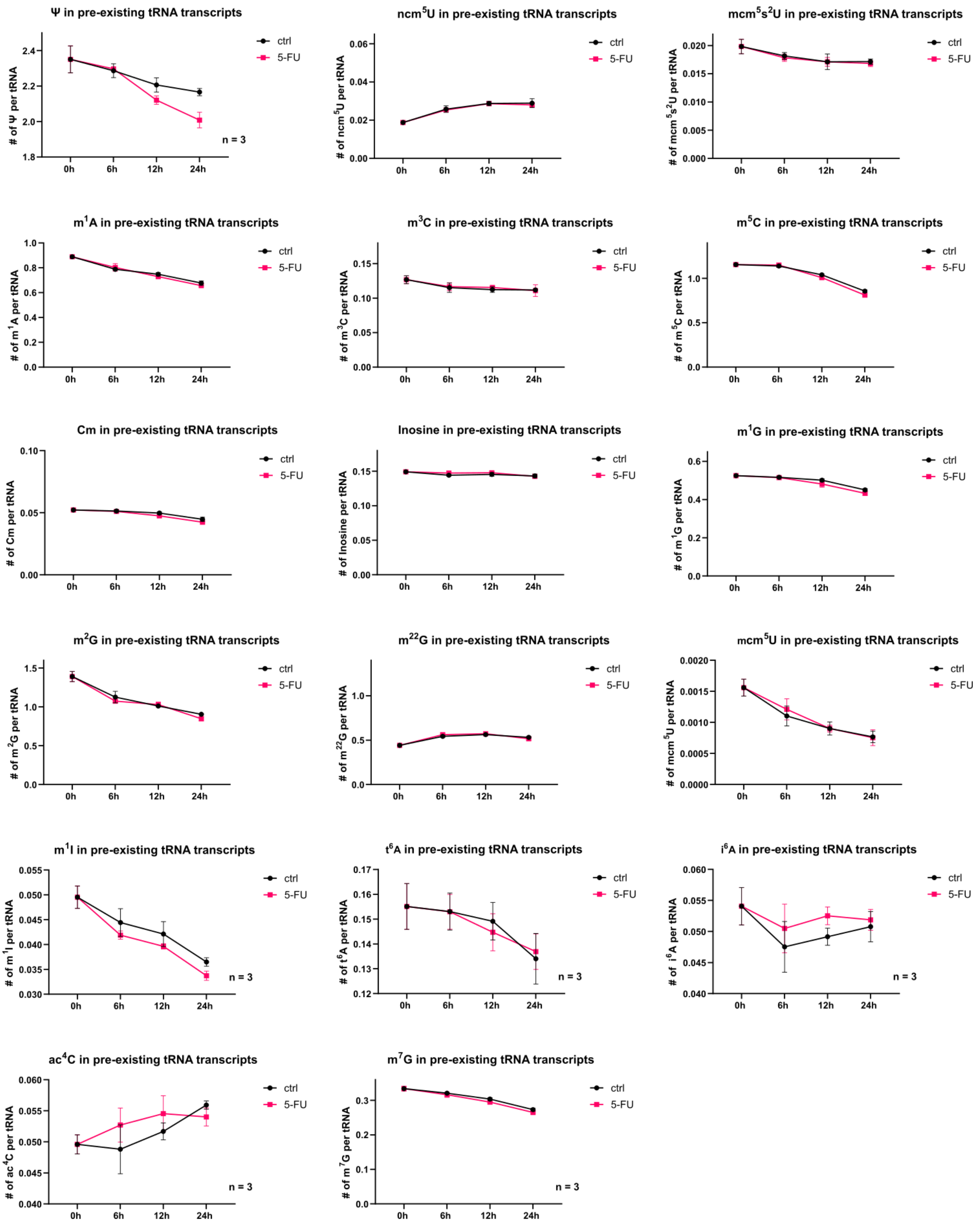


Figure S5.2: Modification profile of HEK293T pre-existing tRNA transcripts during 5-FU exposure. Cells were harvested after 6h, 12h and 24h, respectively. Modifications in pre-existing tRNA transcripts were normalized using the molar amount of labeled, modified nucleoside divided by the molar amount of labeled tRNA. Labeled tRNA molar amount was calculated using the labeled, canonical nucleosides.

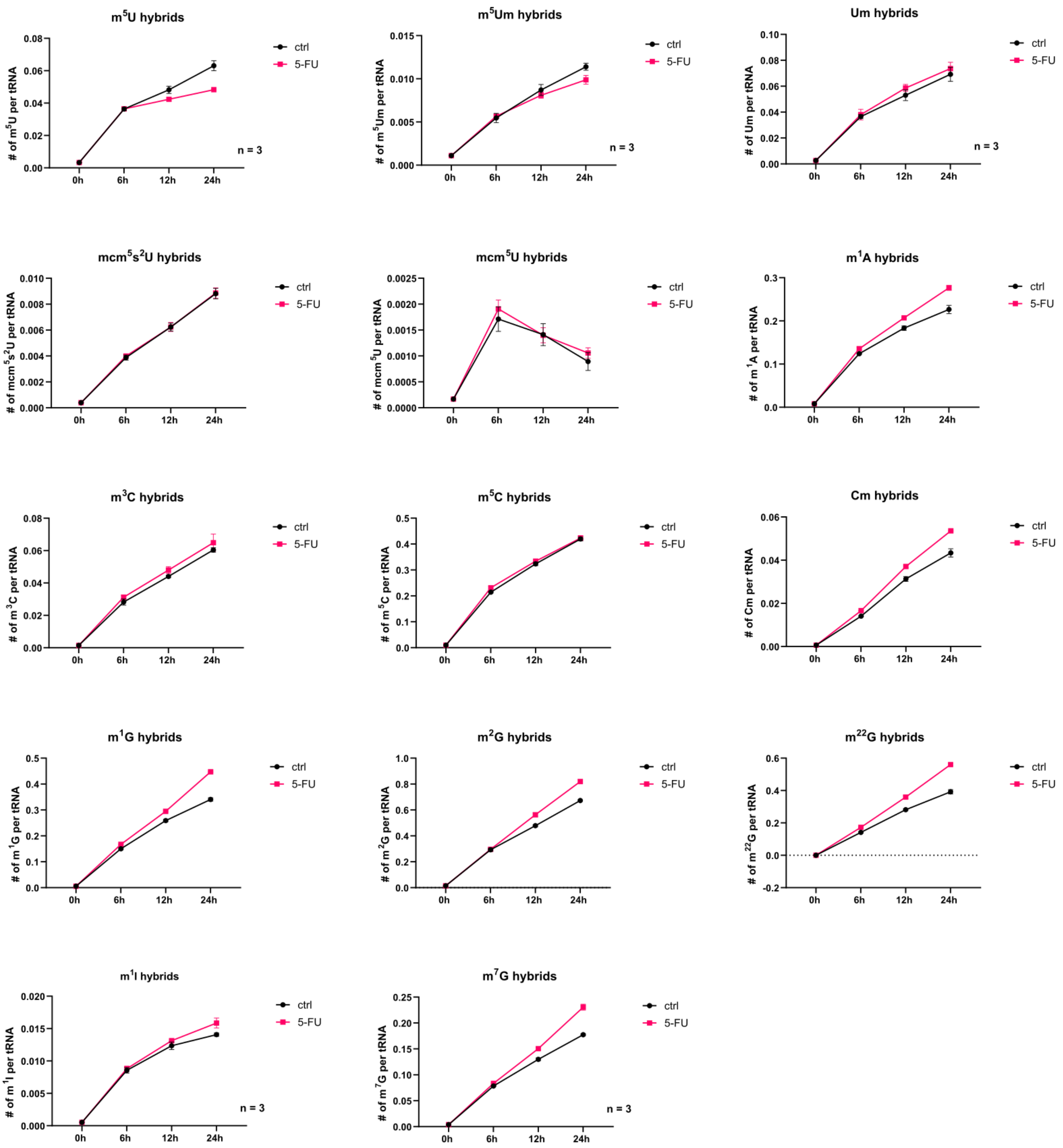


Figure S5.3: Modification profile of HEK293T hybrid tRNA transcripts during 5-FU exposure. Cells were harvested after 6h, 12h and 24h, respectively. Hybrid modifications consist of a labeled nucleoside core-structure and an unlabeled methylgroup (no $CD_3$-labeling). Hybrid modifications were normalized using the molar amount of hybrid, modified nucleoside divided by the molar amount of labeled tRNA. Labeled tRNA molar amount was calculated using the labeled, canonical nucleosides.

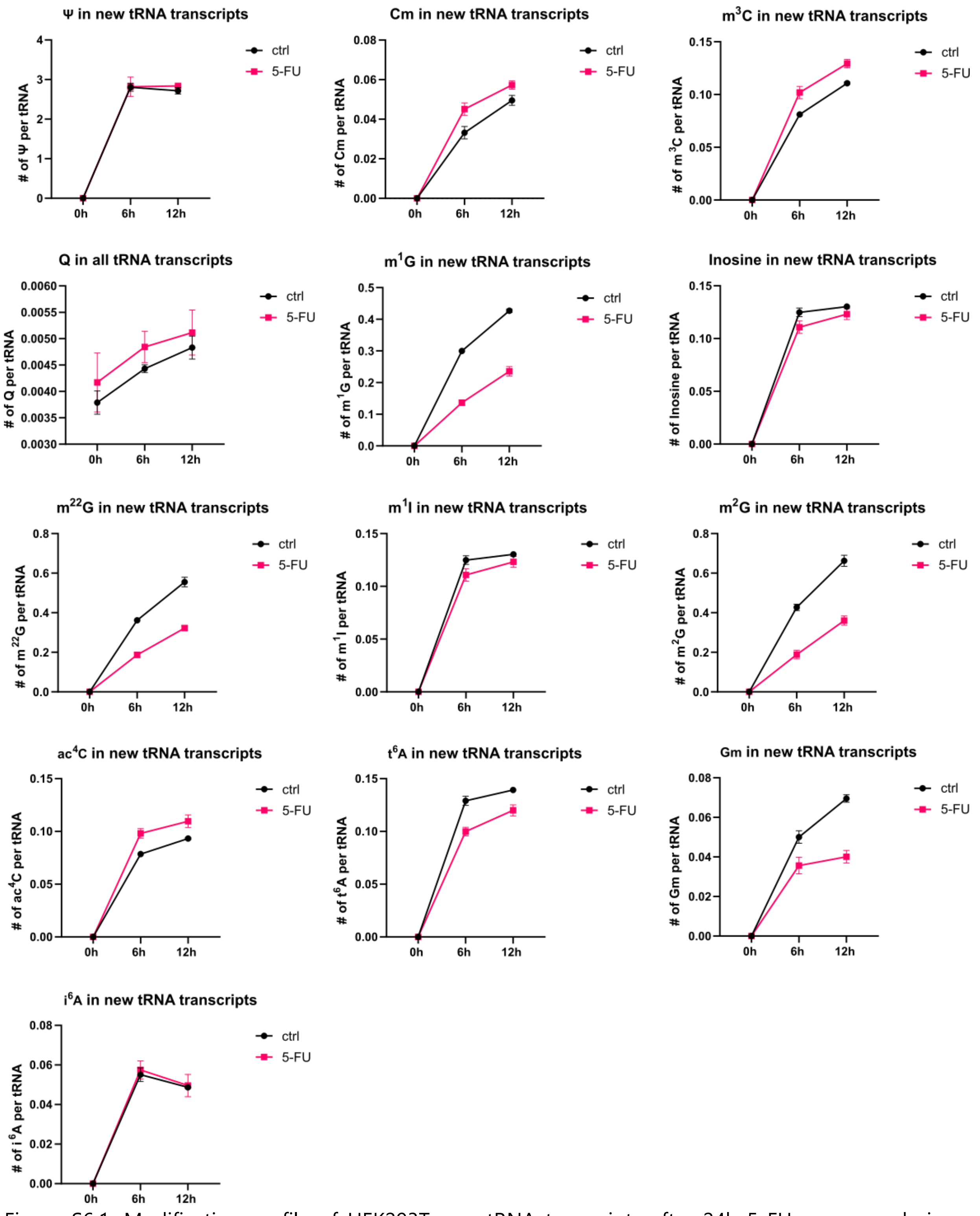


Figure S6.1: Modification profile of HEK293T new tRNA transcripts after 24h 5-FU exposure during recovery (wash-out). Cells were harvested after 6h and 12h, respectively. Modifications in new tRNA transcripts were normalized using the molar amount of unlabeled, modified nucleoside divided by the molar amount of unlabeled tRNA. Unlabeled tRNA molar amount was calculated using the unlabeled, canonical nucleosides.

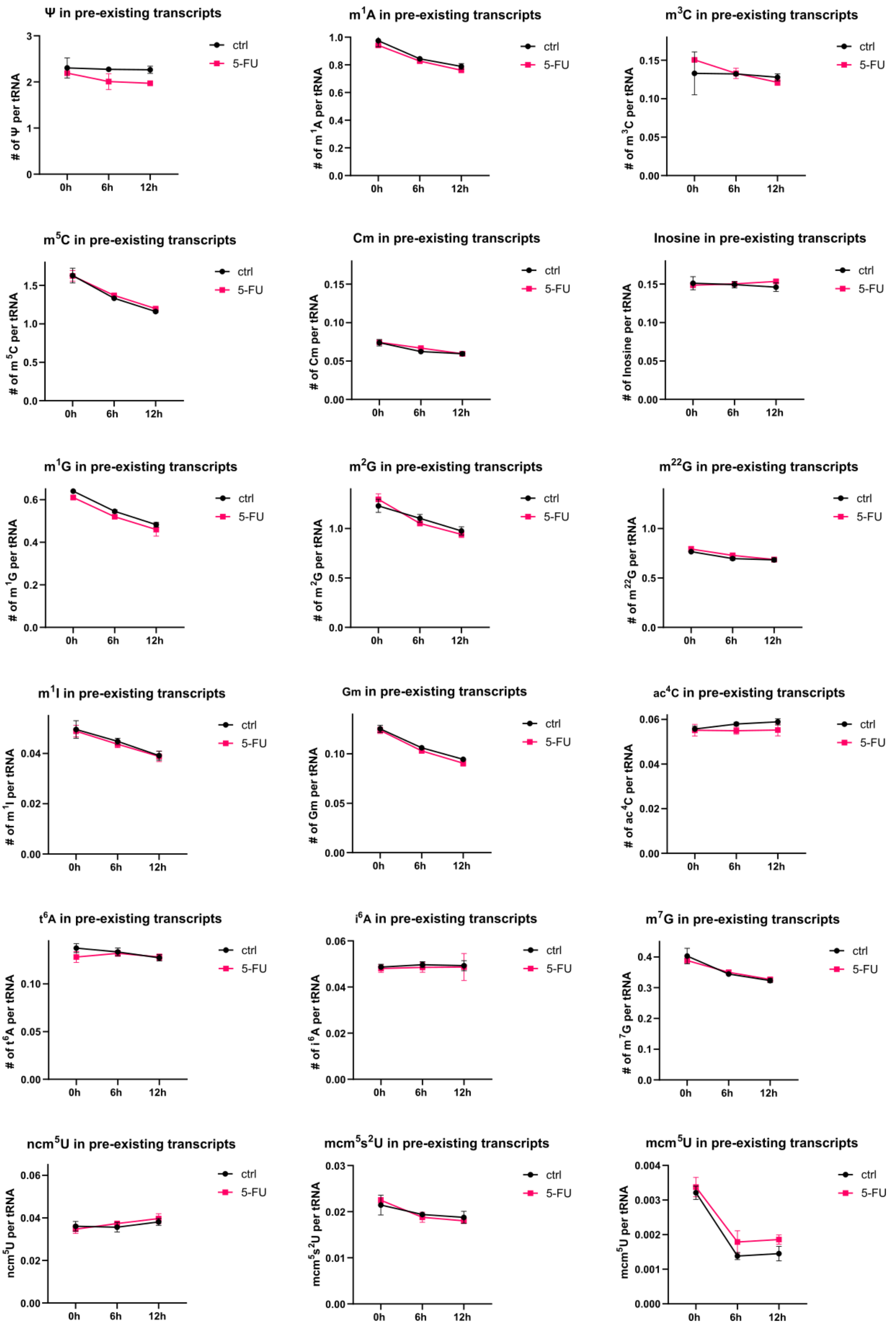


Figure S6.2: Modification profile of HEK293T pre-existing tRNA transcripts after 24h 5-FU exposure during recovery (wash-out). Cells were harvested after 6h and 12h, respectively. Modifications in pre-existing tRNA transcripts were normalized using the molar amount of labeled, modified nucleoside divided by the molar amount of labeled tRNA. Labeled tRNA molar amount was calculated using the labeled, canonical nucleosides.

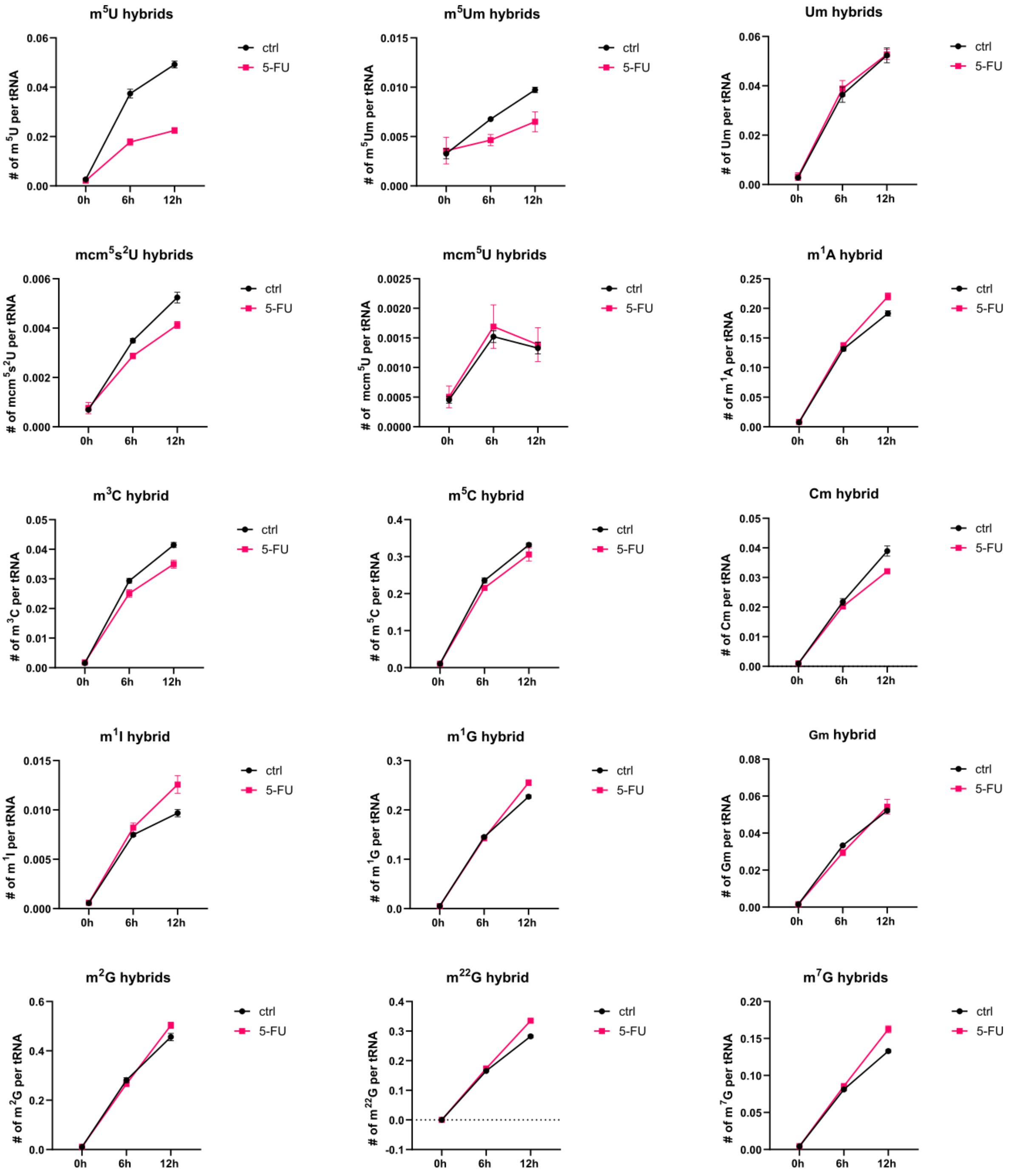


Figure S6.3: Modification profile of HEK293T hybrid tRNA transcripts after 24h 5-FU exposure during recovery (wash-out). Cells were harvested after 6h and 12h, respectively. Hybrid modifications consist of a labeled nucleoside core-structure and an unlabeled methylgroup (no $CD_3$-labeling). Hybrid modifications were normalized using the molar amount of hybrid, modified nucleoside divided by the molar amount of labeled tRNA. Labeled tRNA molar amount was calculated using the labeled, canonical nucleosides.

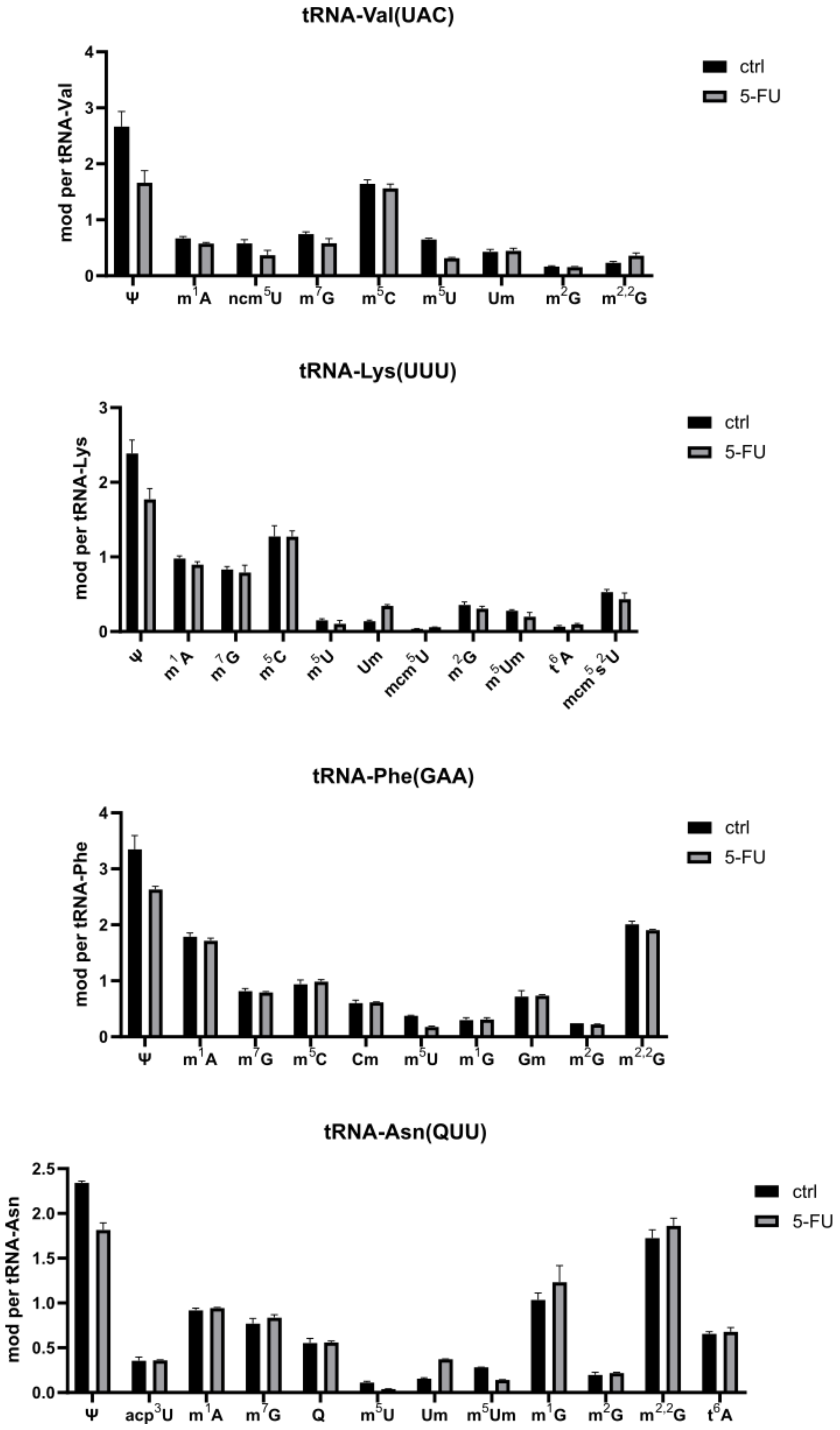


Figure S7.1: Summary of HEK293T modification profile for tRNA-Val-UAC, tRNA-Lys-UUU, tRNA-Phe-GAA and tRNA-Asn-QUU after 24h 5-FU exposure.

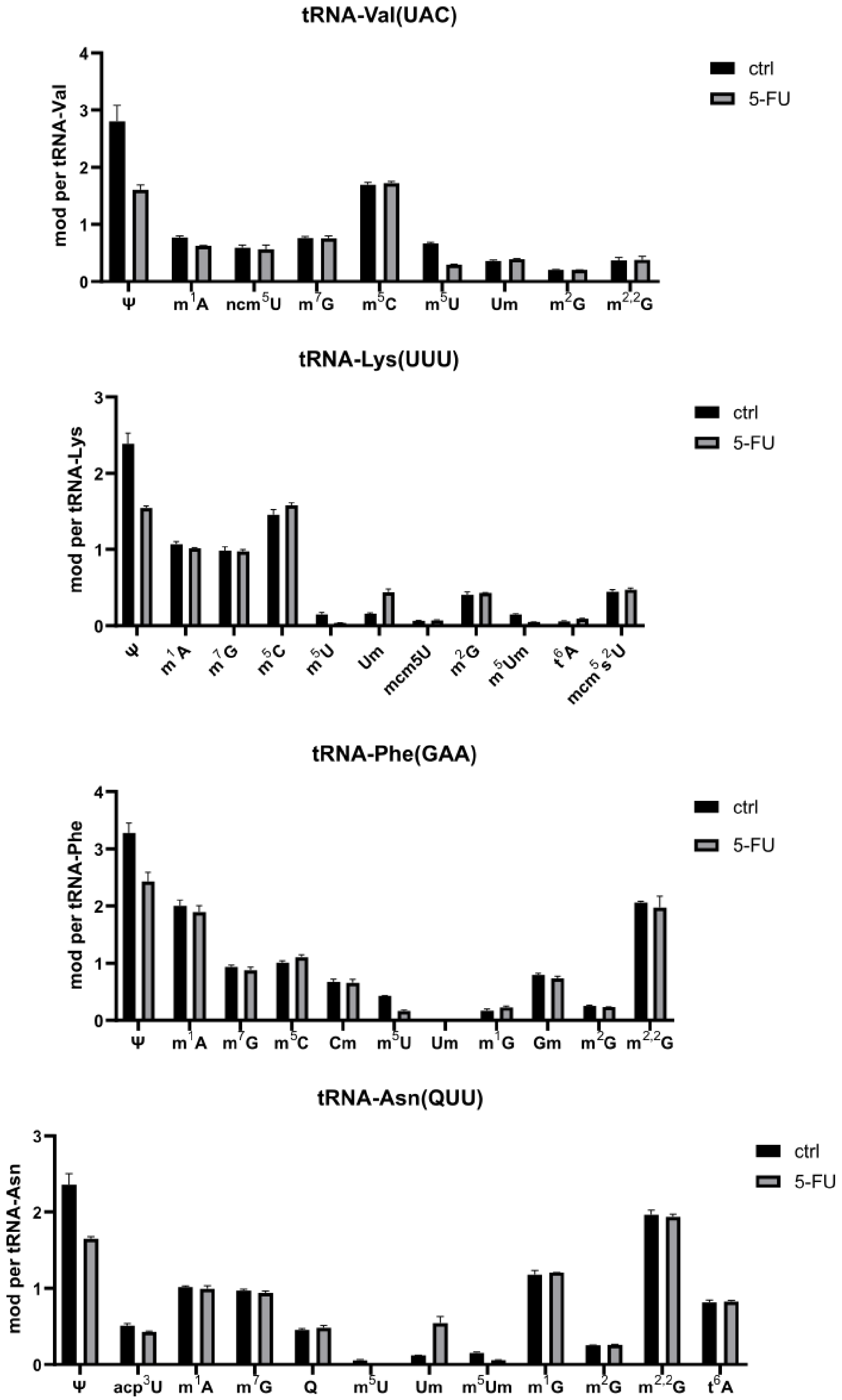


Figure S7.2: Summary of HEK293T modification profile for tRNA-Val-UAC, tRNA-Lys-UUU, tRNA-Phe-GAA and tRNA-Asn-QUU after 24h 5-FU exposure and a 6h recovery duration.

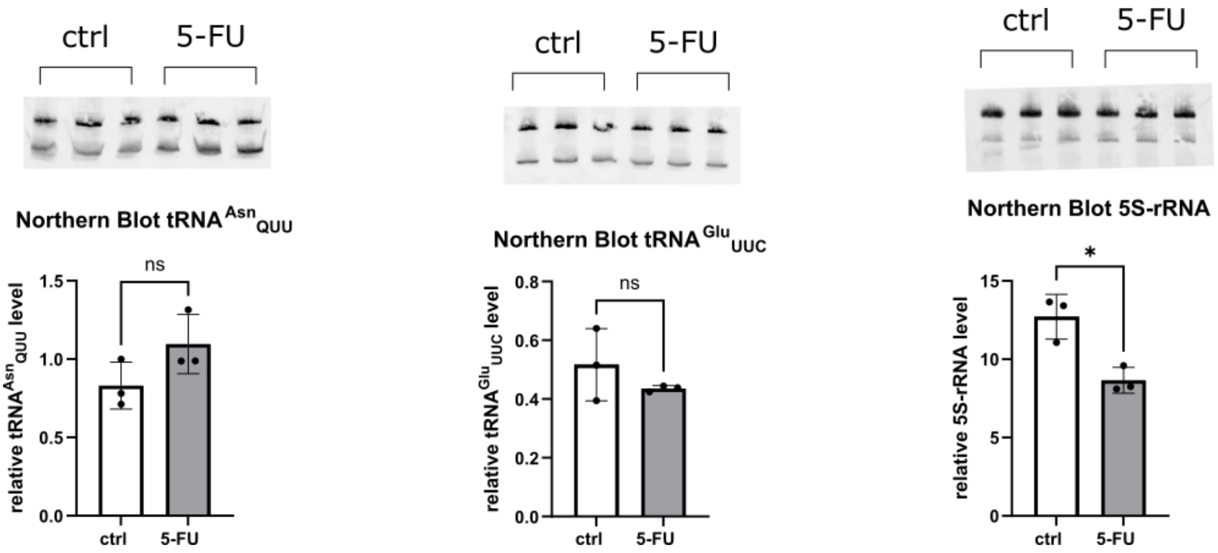


Figure S8: Northern Blotting of tRNA-Asn-QUU, tRNA-Glu-UUC and 5S-rRNA after 24h 5-FU exposure. Relative isoacceptor/rRNA-level was calculated using U6-snRNA as a reference. Detection method: Cy3-labeled oligonucleotides, reverse-complementary to RNA of interest.

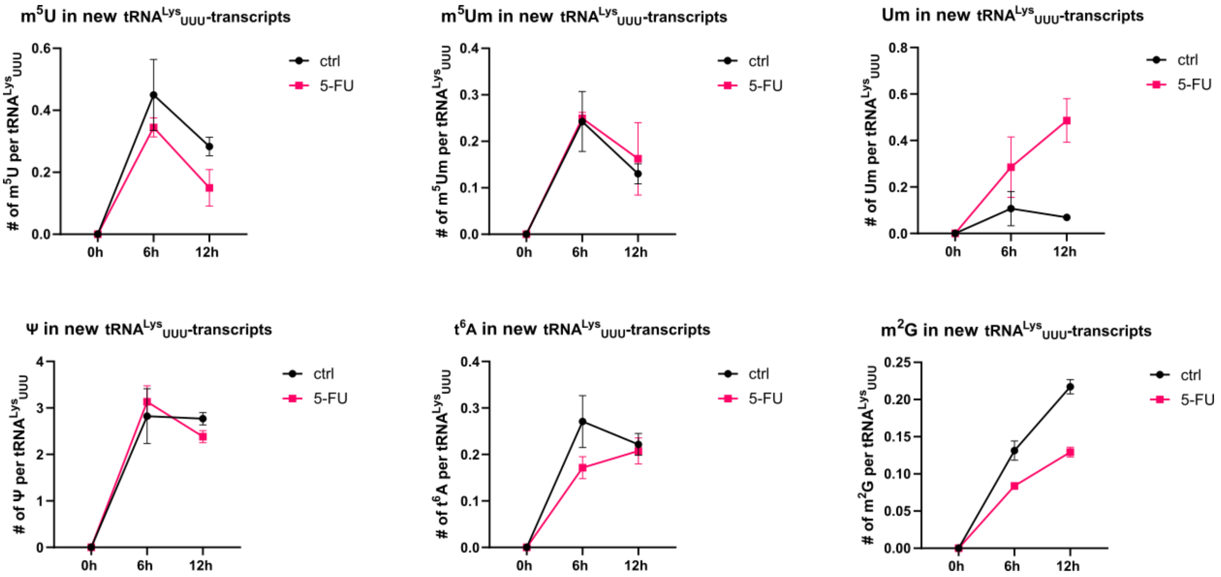


Figure S9.1: Modification profile of HEK293T new $tRNA^{Lys}_{UUU}$-transcripts after 24h 5-FU treatment during recovery (wash-out). Cells were harvested after 6h and 12h, respectively. Modifications in new tRNA transcripts were normalized using the molar amount of unlabeled, modified nucleoside divided by the molar amount of unlabeled $tRNA^{Lys}_{UUU}$. Unlabeled $tRNA^{Lys}_{UUU}$ molar amount was calculated using the unlabeled, canonical nucleosides.

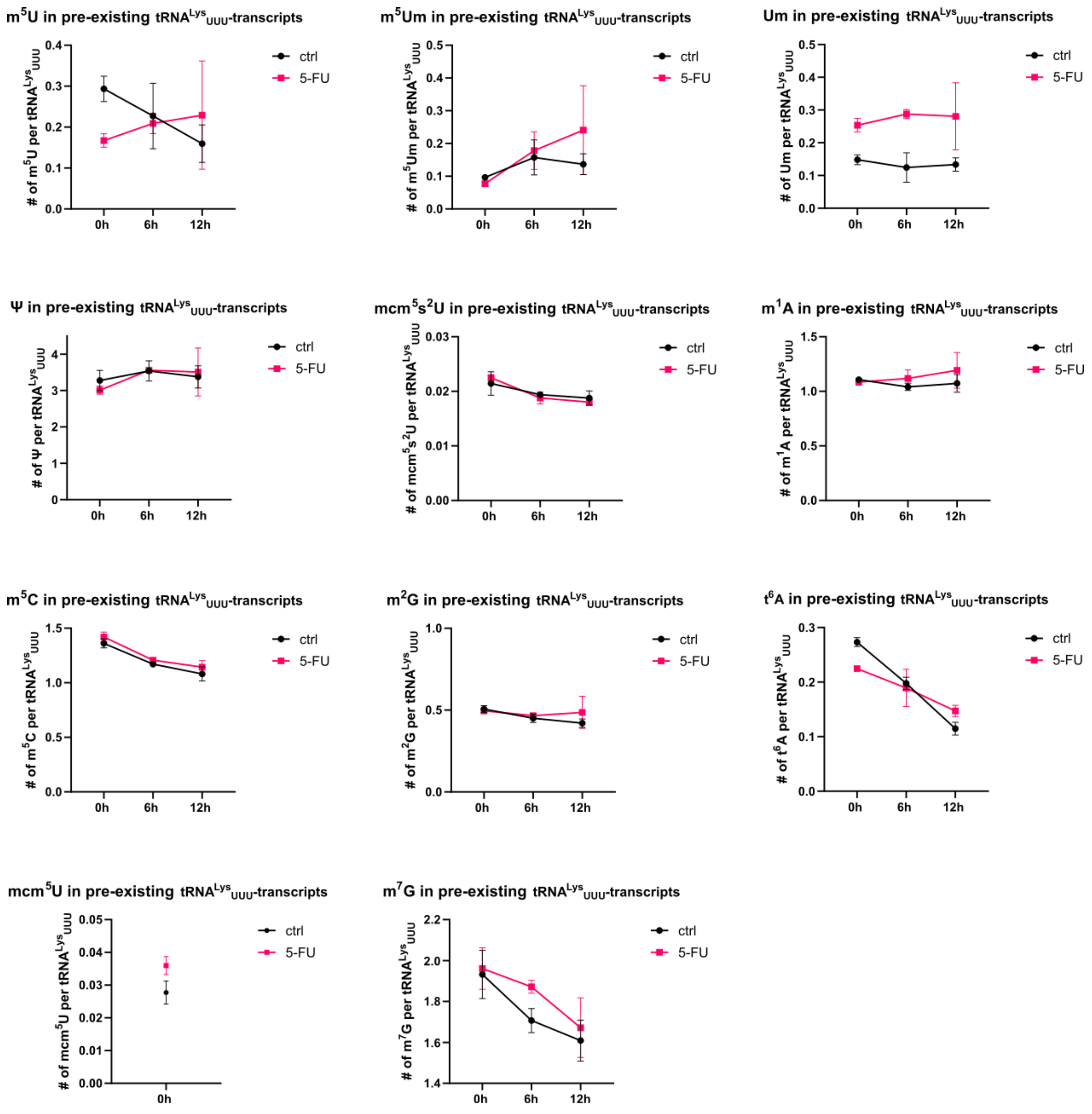


Figure S9.2: Modification profile of HEK293T pre-existing tRNA$^{Lys}_{UUU}$-transcripts after 24h 5-FU treatment during recovery (wash-out). Cells were harvested after 6h and 12h, respectively. Modifications in pre-existing tRNA transcripts were normalized using the molar amount of labeled, modified nucleoside divided by the molar amount of labeled tRNA$^{Lys}_{UUU}$. Labeled tRNA$^{Lys}_{UUU}$ molar amount was calculated using the labeled, canonical nucleosides.

**Note: Hybrid-transcript results are not shown for tRNA-Lys-UUU due to low signals.**

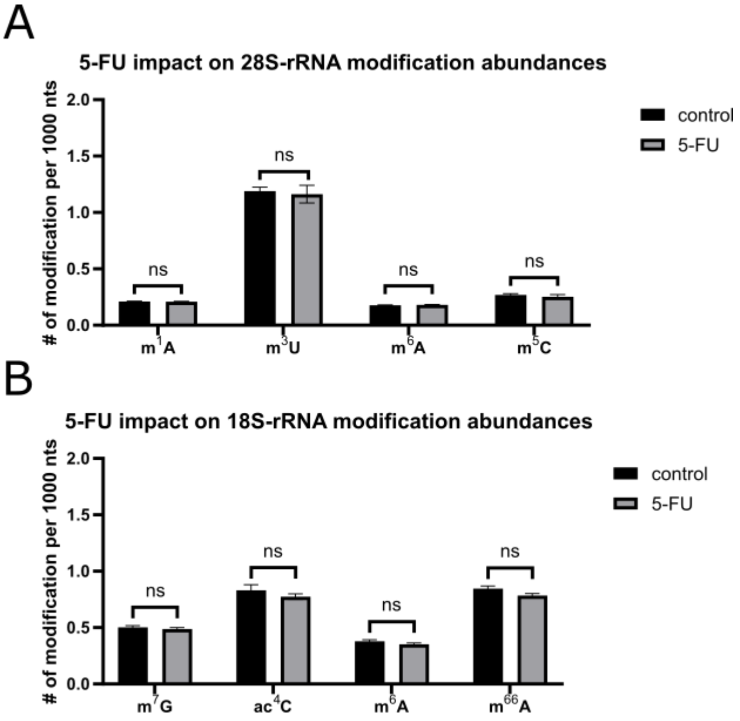


Figure S10: Impact of 5-FU on rRNA modification abundances. **(A)** 28S-rRNA modification abundances for modifications commonly found in this subtype after 24h 5-FU exposure. The number of modification was calculated using the molar amount of modified nucleoside normalized on the molar amount of 1000 canonical nucleosides. **(B)** 18S-rRNA modification abundances for modifications commonly found in this subtype after 24h 5-FU exposure. The number of modification was calculated using the molar amount of modified nucleoside normalized on the molar amount of 1000 canonical nucleosides.

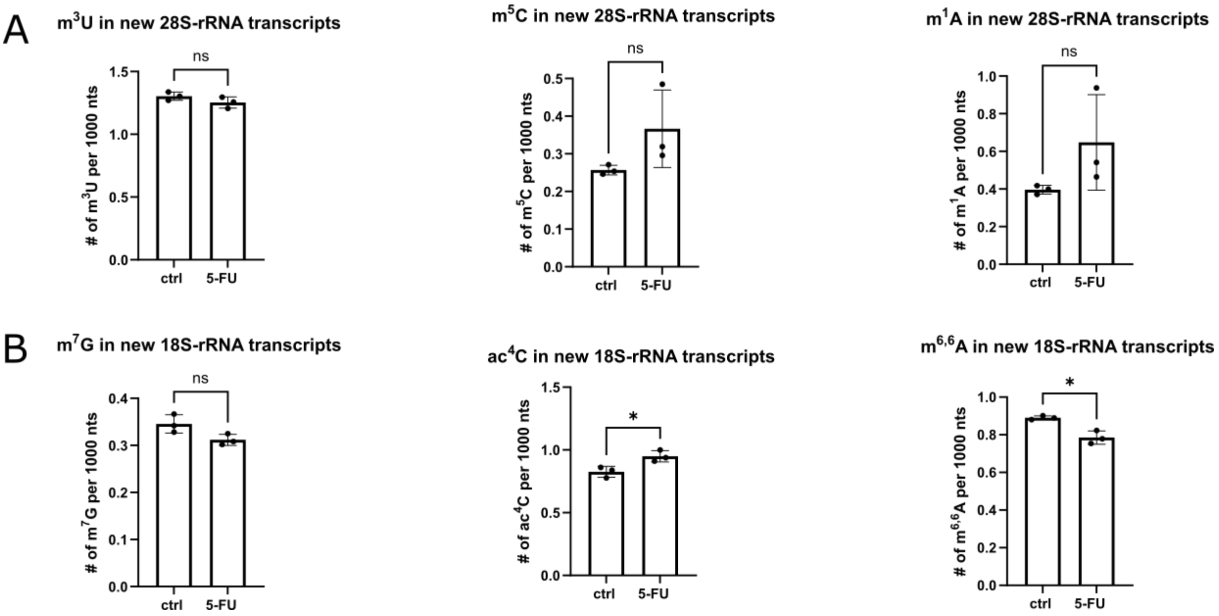


Figure S11.1: Modification abundances in new rRNA transcripts after 24h 5-FU exposure using NAIL-MS. **(A)** Modification abundances in new 28S-rRNA transcripts. The number of modification was referenced on the molar amount of 1000 canonical nucleosides. **(B)** Modification abundances in new 18S-rRNA transcripts. The number of modification was referenced on the molar amount of 1000 canonical nucleosides.

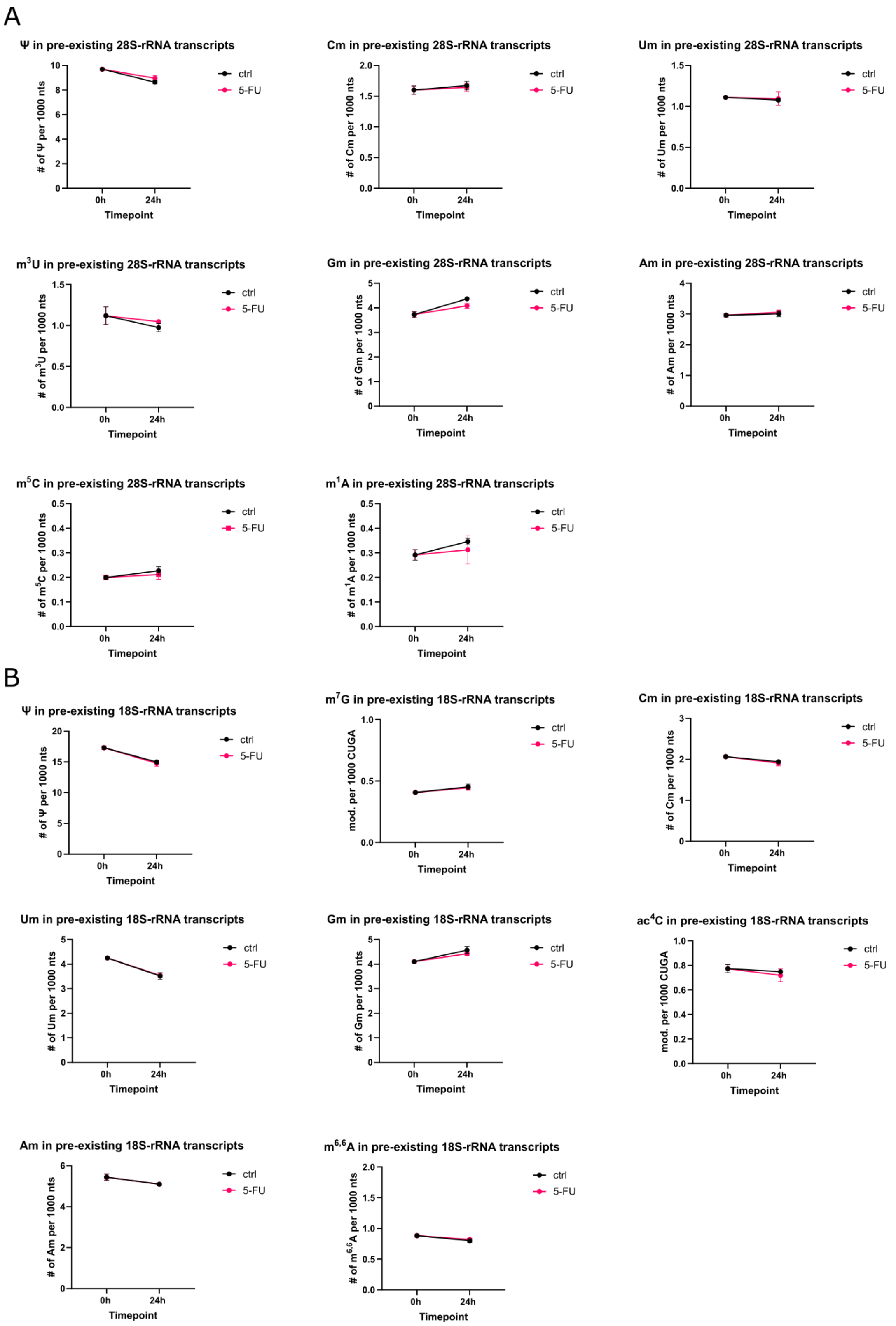


Figure S11.2: Modification abundances in pre-existing rRNA transcripts during 5-FU exposure using NAIL-MS. **(A)** Modification abundances in pre-existing 28S-rRNA transcripts. Cells were harvested after 24h of treatment. The number of modification was referenced on the molar amount of 1000 canonical nucleosides. **(B)** Modification abundances in pre-existing 18S-rRNA transcripts. Cells were harvested after 24h of treatment. The number of modification was referenced on the molar amount of 1000 canonical nucleosides. Note: Control cells before experiment initiation were harvested (t = 0 h).

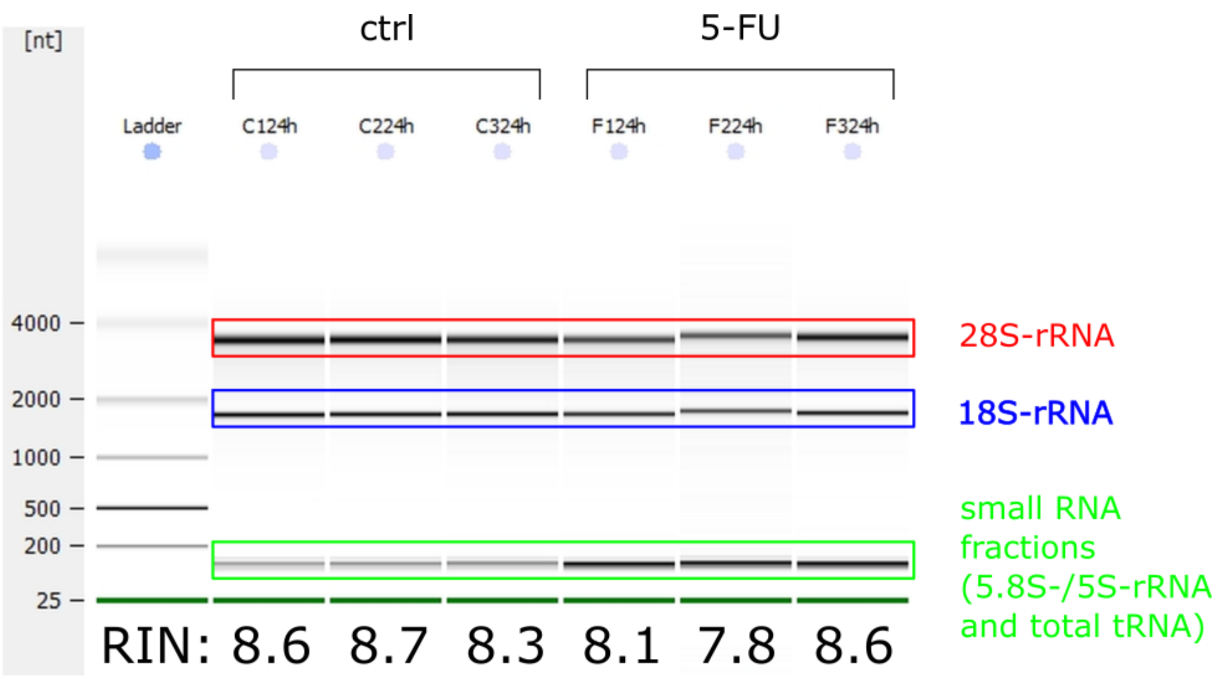


Figure S12: Bioanalyzer pico chip for HEK293T total RNA integrity check (quality control). Cells were harvested after 24h 5-FU exposure. Ladder shows respective RNA length on the left. Bands are highlighted for 28S-rRNA (red), 18S-rRNA (blue) and the combined small RNA fraction (green). RNA integrity number (RIN) is shown below the respective lane.

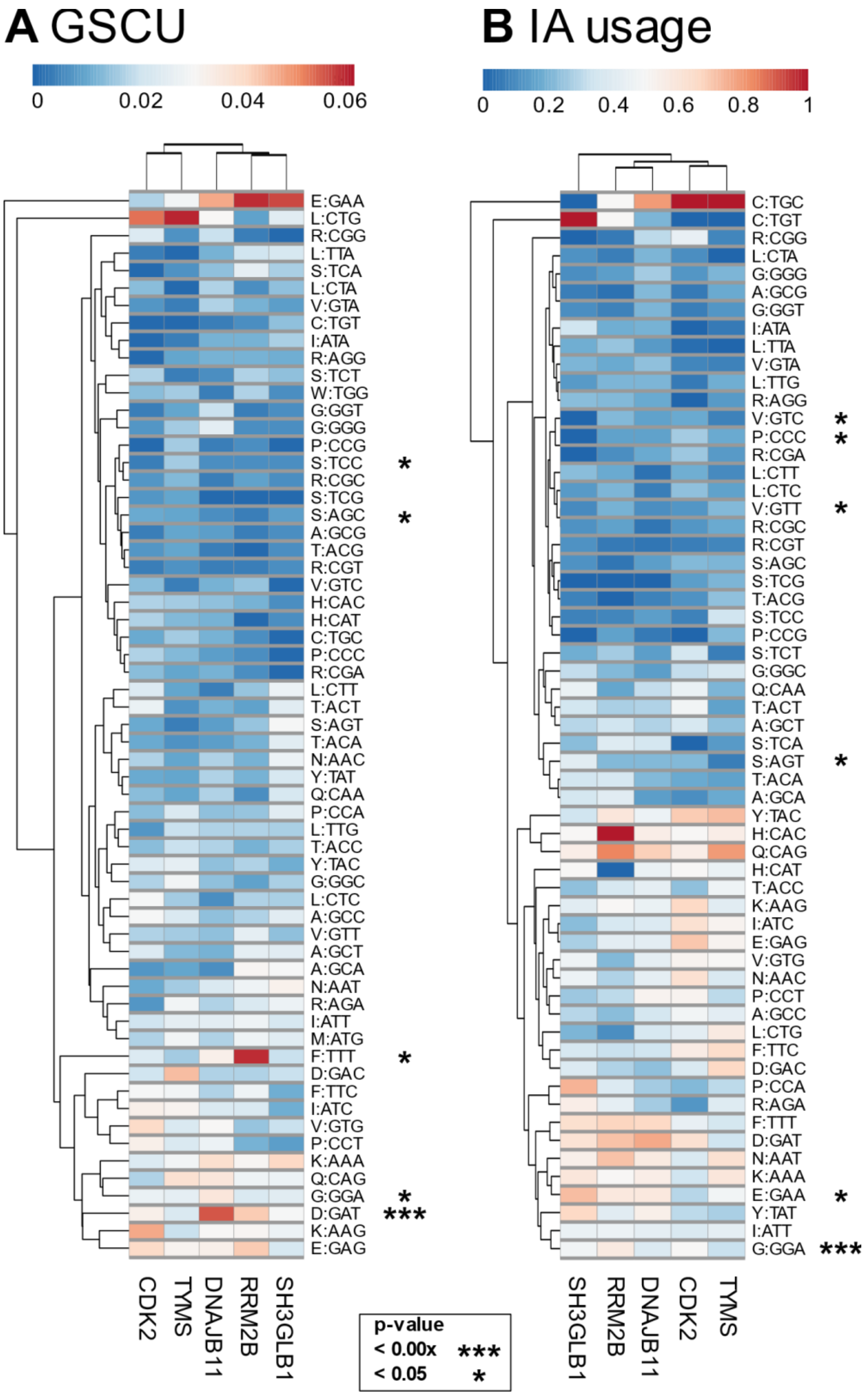


Figure S13: Codon usage analysis of proteins highly expressed after 24 hours of 5-FU treatment. **A** Gene specific codon usage analysis (GSCU) of 5 upregulated genes. **B** Isoacceptor usage of 5 upregulated genes. Statistics: t-test of 5 upregulated genes against whole genome. p-values are indicated as stars. Clustered using clustvis (1) using Euclidean distance and column average clustering method.

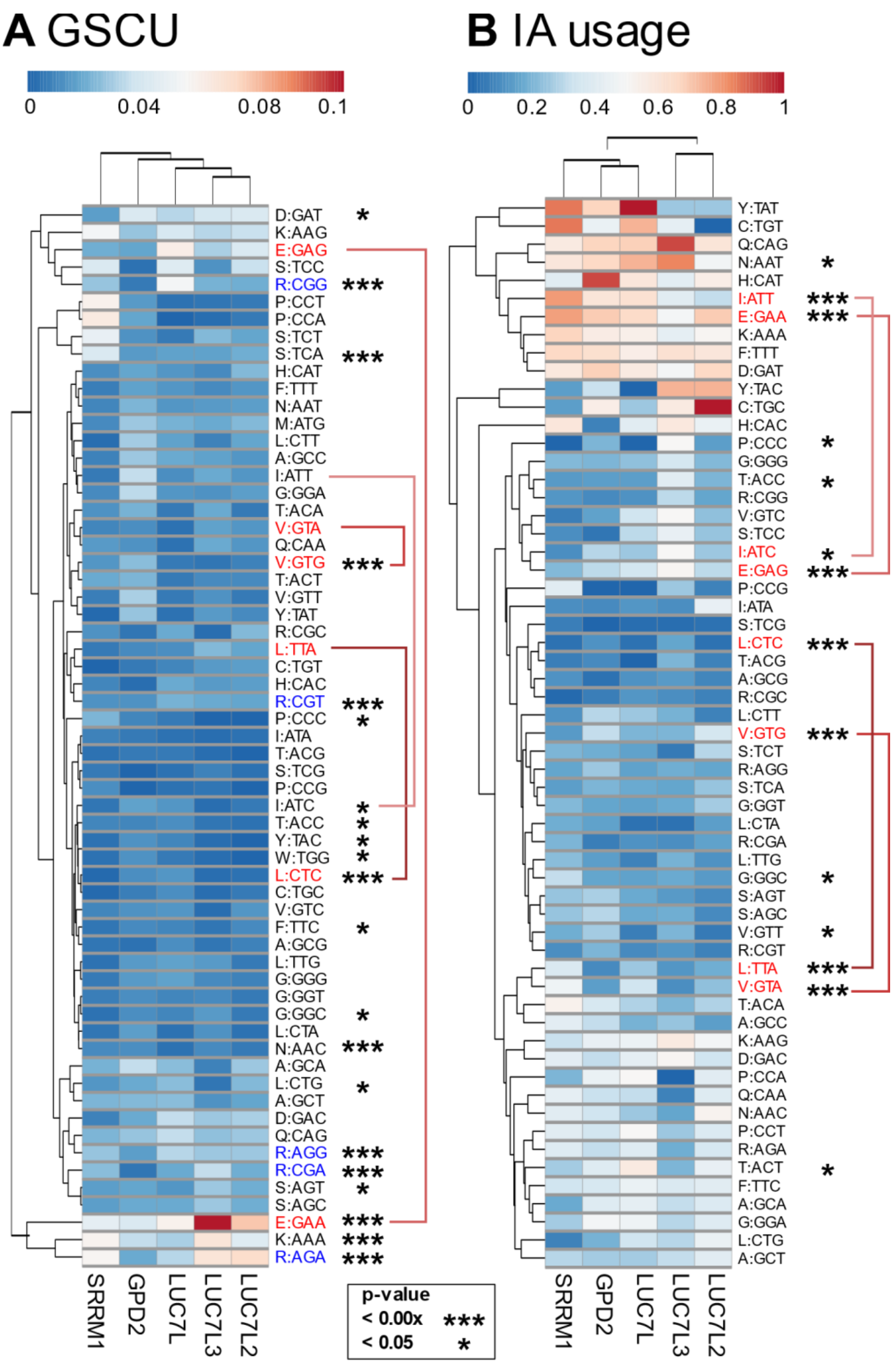


Figure S14: Codon usage analysis of proteins highly expressed after 6 hours recovery following 24 hours of 5-FU treatment. Note: Analysis of LYPLA1 and GPD2 was not possible, genes are missing in the database. **A** Gene specific codon usage analysis (GSCU) of 5 upregulated genes. **B** Isoacceptor usage of 5 upregulated genes. Statistics: t-test of 5 upregulated genes against whole genome. p-values are indicated as stars. Clustered using clustvis (1) using Euclidean distance and column average clustering method. Square brackets connect significantly changed codons of same amino acid.

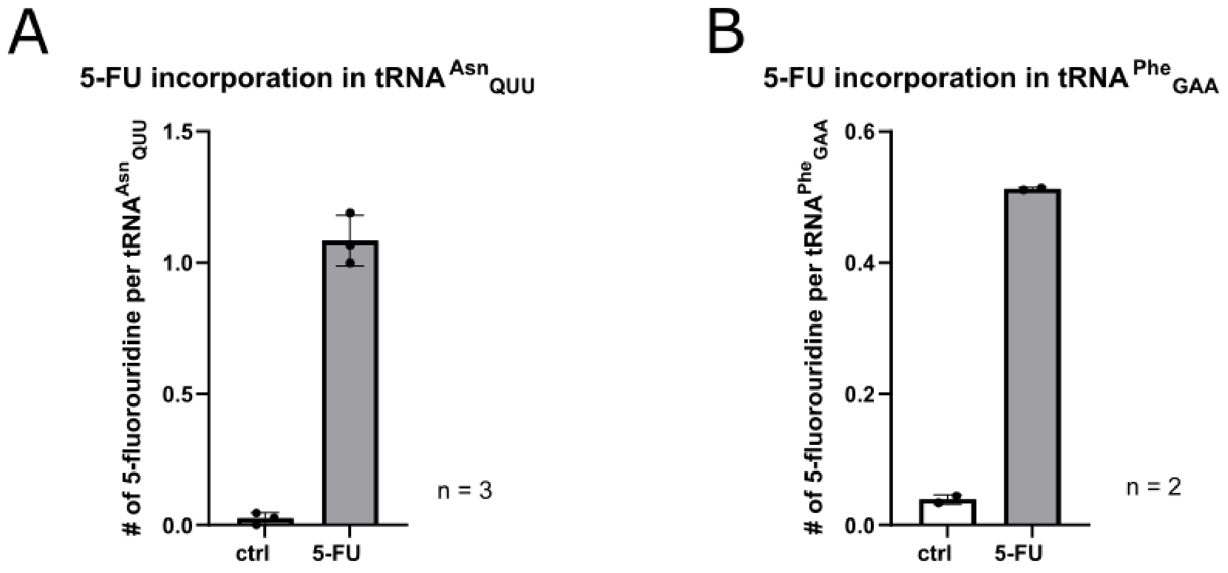


Figure S15: 5-FU incorporation as 5-fluorouridine in individual isoacceptors obtained by isoacceptor purification protocol (as described in the materials methods part of the manuscript) following absolute quantification using nucleoside LC-MS **(A)** 5-fluorouridine level in $tRNA^{Asn}_{QUU}$ after 24h of acute 5-FU exposure following 6h of recovery. **(B)** 5-fluorouridine level in $tRNA^{Phe}_{GAA}$ after 24h of acute 5-FU exposure.

| Table S1: Proteins involved in tRNA metabolism detected in 5-FU acute exposure shotgun-proteomics | | | |
|---|---|---|---|
| log p-value | Difference | Protein names | Gene names |
| 0,95977642 | -0,22241936 | Histidine--tRNA ligase, cytoplasmic | HARS |
| 0,04042513 | 0,03133774 | tRNA pseudouridine synthase;tRNA pseudouridine synthase A, mitochondrial | PUS1 |
| 1,03108447 | -0,30123405 | Asparagine--tRNA ligase, cytoplasmic | NARS |
| 1,63210769 | -0,1143837 | Bifunctional glutamate/proline--tRNA ligase;Glutamate--tRNA ligase;Proline--tRNA ligase | EPRS |
| 0,83083548 | -0,19040489 | Aspartate--tRNA ligase, cytoplasmic | DARS |
| 0,69928738 | -0,14869843 | Tryptophan--tRNA ligase, cytoplasmic;T1-TrpRS;T2-TrpRS | WARS |
| 0,83231392 | -0,23000298 | Threonine--tRNA ligase, cytoplasmic | TARS |
| 0,60263785 | -0,14020233 | Valine--tRNA ligase | VARS |
| 0,8731757 | 0,37604065 | Glycine--tRNA ligase | GARS |
| 0,59659675 | -0,0841404 | Isoleucine--tRNA ligase, cytoplasmic | IARS |
| 0,56315691 | -0,11359215 | Glutamine--tRNA ligase | QARS |
| 0,41134861 | -0,08953705 | Alanine--tRNA ligase, cytoplasmic | AARS |
| 0,95807583 | -0,10387154 | Serine--tRNA ligase, cytoplasmic | SARS |
| 1,07426135 | -0,17658005 | Arginine--tRNA ligase, cytoplasmic | RARS |
| 0,18890307 | -0,0432003 | Tyrosine--tRNA ligase, cytoplasmic;Tyrosine--tRNA ligase, cytoplasmic, N-terminally processed;Tyrosine--tRNA ligase | YARS |
| 0,56083762 | -0,10071144 | Methionine--tRNA ligase, cytoplasmic | MARS |
| 0,35565594 | -0,07700996 | tRNA (cytosine(34)-C(5))-methyltransferase | NSUN2 |
| 0,53844818 | -0,15674591 | Lysine--tRNA ligase | KARS |
| 0,19163896 | 0,11092834 | Pseudouridylate synthase 7 homolog | PUS7 |
| 0,54528362 | 0,21651688 | Elongator complex protein 3 | ELP3 |
| 2,22172082 | -0,380513 | Phenylalanine--tRNA ligase beta subunit | FARSB |
| 0,47193759 | -0,10910378 | Isoleucine--tRNA ligase, mitochondrial | IARS2 |
| 0,21635543 | 0,18072166 | tRNA (guanine(26)-N(2))-dimethyltransferase | TRMT1 |
| 1,41544699 | -0,32334213 | Leucine--tRNA ligase, cytoplasmic | LARS |
| 0,25125783 | -0,15986748 | tRNA (guanine-N(7)-)-methyltransferase | METTL1 |
| 0,3251369 | 0,11245079 | Multifunctional methyltransferase subunit TRM112-like protein | TRMT112 |
| 1,4741189 | -0,33277206 | Phenylalanine--tRNA ligase alpha subunit | FARSA |
| 1,77951304 | -0,25374374 | tRNA-splicing ligase RtcB homolog | RTCB |

| Table S2: Proteins involved in tRNA metabolism detected in 5-FU recovery exposure shotgun-proteomics | | | |
|---|---|---|---|
| log p-value | Difference | Protein names | Gene names |
| 0,96959213 | 0,4521904 | tRNA pseudouridine synthase;tRNA pseudouridine synthase A, mitochondrial | PUS1 |
| 0,37028971 | -0,31557503 | tRNA (guanine-N(7)-)-methyltransferase | METTL1 |
| 0,17879017 | 0,08876076 | Peptidyl-tRNA hydrolase 2, mitochondrial | PTRH2 |
| 0,73328566 | 0,12616043 | Asparagine--tRNA ligase, cytoplasmic | NARS |
| 0,26313019 | 0,06088867 | Bifunctional glutamate/proline--tRNA ligase;Glutamate--tRNA ligase;Proline--tRNA ligase | EPRS |
| 0,37807263 | -0,07453651 | Histidine--tRNA ligase, cytoplasmic | HARS |
| 0,05585443 | 0,01255569 | Aspartate--tRNA ligase, cytoplasmic | DARS |
| 0,0963421 | -0,03755608 | Tryptophan--tRNA ligase, cytoplasmic;T1-TrpRS;T2-TrpRS | WARS |
| 0,40422157 | -0,2245842 | Threonine--tRNA ligase, cytoplasmic | TARS |
| 0,5272207 | 0,08509178 | Valine--tRNA ligase | VARS |
| 0,13242388 | -0,14681358 | Glycine--tRNA ligase | GARS |
| 0,08698738 | 0,02595978 | Isoleucine--tRNA ligase, cytoplasmic | IARS |
| 0,04114511 | 0,01710548 | Glutamine--tRNA ligase | QARS |
| 2,3731635 | 0,28981972 | Alanine--tRNA ligase, cytoplasmic | AARS |
| 0,297843 | 0,06765213 | Serine--tRNA ligase, cytoplasmic | SARS |
| 0,18433666 | -0,05220222 | Arginine--tRNA ligase, cytoplasmic | RARS |
| 0,02072048 | -0,01045685 | Tyrosine--tRNA ligase, cytoplasmic;Tyrosine--tRNA ligase, cytoplasmic, N-terminally processed;Tyrosine--tRNA ligase | YARS |
| 0,58297495 | 0,21817856 | Methionine--tRNA ligase, cytoplasmic | MARS |
| 0,66175178 | 0,16958199 | tRNA (cytosine(34)-C(5))-methyltransferase | NSUN2 |
| 0,56867953 | 0,11972198 | Aminoacyl tRNA synthase complex-interacting multifunctional protein 1;Endothelial monocyte-activating polypeptide 2 | AIMP1 |
| 1,39145323 | -0,36834526 | Aminoacyl tRNA synthase complex-interacting multifunctional protein 2 | AIMP2 |
| 0,7254734 | 0,15158119 | Lysine--tRNA ligase | KARS |
| 0,11850261 | -0,1455368 | Putative peptidyl-tRNA hydrolase PTRHD1 | PTRHD1 |
| 0,10637032 | -0,11385193 | Alanine--tRNA ligase, mitochondrial | AARS2 |
| 0,45194307 | -0,34320946 | Aspartate--tRNA ligase, mitochondrial | DARS2 |
| 0,11646487 | -0,09310608 | Queuine tRNA-ribosyltransferase subunit QTRTD1 | QTRTD1 |
| 1,33530306 | -0,20408592 | Phenylalanine--tRNA ligase beta subunit | FARSB |
| 0,12588353 | -0,04406929 | Isoleucine--tRNA ligase, mitochondrial | IARS2 |
| 0,04876499 | 0,05223541 | tRNA (guanine(26)-N(2))-dimethyltransferase | TRMT1 |
| 0,53560902 | -0,19715767 | Leucine--tRNA ligase, cytoplasmic | LARS |
| 0,66662055 | 0,19273567 | Phenylalanine--tRNA ligase alpha subunit | FARSA |
| 1,56966689 | -0,16431313 | tRNA-splicing ligase RtcB homolog | RTCB |

| | | | |
|---|---|---|---|
| 1,09864268 | 0,36981201 | Multifunctional methyltransferase subunit TRM112-like protein | TRMT112 |
| 0,65769058 | -0,44917717 | Pseudouridylate synthase 7 homolog | PUS7 |
| 0,17363559 | 0,13048935 | Elongator complex protein 3 | ELP3 |

Table S3: Oligonucleotide sequences for Northern Blotting and tRNA-isoacceptor purification.

| tRNA-isoacceptor [purpose] | Oligonucleotide sequence incl. tags [Btn = Biotin; Cy3 = Cyanine 3] |
|---|---|
| tRNA-Phe(GAA) [IA purification] | [Btn]AAATGGTGCCGAAACCCGGGATCGAACCAGGGT |
| tRNA-Val(UAC) [IA purification] | [Btn]GGTTCCACTGGGGCTCGAACCCAGGACCTTCTGCGT |
| tRNA-Lys(UUU) [IA purification] | GAACCCTGGACCCTCAGATTAAAAGTAAA[BtnTg] |
| tRNA-Asn(QUU) [IA purification] | AACCACCAACCTTTCGGTTAACAGCCAAA[BtnTg] |
| tRNA-Phe(GAA) [Northern Blotting] | [Cyanine3]TGGTGCCGAAACCCGGGATCGAACCAGGGT[Cyanine3] |
| tRNA-Val(UAC) [Northern Blotting] | [Cyanine3]CGTTCCACTGGGGCTCGAACCCAGGACCTTCTGCGT[Cyanine3] |
| tRNA-Lys(UUU) [Northern Blotting] | [Cyanine3]TGGCGCCCGAACAGGGACTTGAACCCTGGACCCTCAGATTAAAAGTCTGATGCTCTACCGACTGAGCTATCCGGGC[Cyanine3] |
| tRNA-Asn(QUU) [Northern Blotting] | [Cyanine3]TGGCGTCCCTGGGTGGGCTCGAACCACCAACCTTTCGGTTAACAGCCGAACGCGCTAACCGATTGCGCCACAGAGAC[Cyanine3] |
| tRNA-Glu(UUC) [Northern Blotting] | [Cyanine3]CCAGGAATCCTAACCGCTAGACCATRTGGA[Cyanine3] |
| 5S-rRNA [Northern Blotting] | [Cyanine3]AAACCGACCCTGCTTAGCTTCCGAGATCAGACG[Cyanine3] |
| U6-snRNA [Northern Blotting] | [Cyanine3]AAATATGGAACGCTTCACGAATTTGCGTGTCATCCTTGC[Cyanine3] |

Table S5: Comparison between Translatome and Proteome data. Given are the changes in protein expression after 5-FU treatment obtained by our proteomics results (data taken from both acute and recovery 5-FU incubation). We then compared the changes in protein expression with changes in the respective actively translated mRNA transcripts obtained by ribosome profiling (also see PMID 35013311). Significant, pairing matches between both datasets are marked in red.

| **Upregulated proteins** | **Downregulated proteins** |
|---|---|
| SRSF11 | RPL32 |
| CDK2 | RPL15 |
| TYMS | RPL13 |
| UQCR1 | RPL30 |
| RRM2 | RPL6 |
| SH3GLB1 | RPL8 |
| DNAJB11 | RPS15 |
| SRRM1 | RPL23A |
| LUC7L | GNL3 |
| PRKACA | RPL3 |
| LYPLA1 | RPL7 |
| GDP2 | RPL4 |
| | RPL13A |
| | RPS4X |
| | BTF3 |
| | RPL10A |
| | RPS17 |
| | RPS7 |
| | EIF1 |
| | IMPDH1 |
| | AAMP |
| | GNB2L1 |
| | RPS24 |
| | RPS12 |
| | CHCHD2 |
| | RPL10 |
| | RPL18 |
| | RPS13 |
| | RPS3A |
| | RPS8 |
| | DYNC1LI1 |
| | RPL7A |
| | RPL27 |
| | RPL5 |
| | RPL23 |
| | ATXN10 |
